# Tumor-targeted hydroxyapatite nanoparticles for dual-mode diagnostic imaging and near-infrared light-triggered photothermal cancer therapy

**DOI:** 10.1101/2025.02.20.639217

**Authors:** Loganathan Palanikumar, Faten Mansoor Yasin, Itgel Munkhjargal, Maylis Boitet, Liaqat Ali, Muhammed Shiraz Ali, Rainer Straubinger, Francisco N. Barrera, Mazin Magzoub

**Affiliations:** Biology Program, Division of Science, New York University Abu Dhabi, Abu Dhabi, United Arab Emirates; Core Technology Platforms, New York University Abu Dhabi, Abu Dhabi, United Arab Emirates; Department of Biochemistry & Cellular and Molecular Biology, University of Tennessee Knoxville, Knoxville, Tennessee, United States

**Keywords:** Cancer, diagnostic imaging, fluorescence, hydroxyapatite, IR-1061 dye, nanoparticles, near-infrared light, photothermal therapy, targeted drug delivery, tumor microenvironment

## Abstract

Photothermal therapy (PTT), which utilizes photothermal agents (PTAs) to induce localized hyperthermia within tumors upon light irradiation, has emerged as a promising cancer treatment strategy. However, low water solubility, poor *in vivo* circulation stability and a lack of tumor specificity of many common PTAs limit their applicability. To address these issues, we have developed a simple, yet highly potent, tumor-targeted nanotheranostic system that consists of lipid/PEG-coated hydroxyapatite nanoparticles (LHAPNs) encapsulating the near-infrared (NIR) photothermal dye IR106 (LHAPNIRs). The lipid coat serves to retain the encapsulated dye and prevent serum protein adsorption and macrophage recognition, which would otherwise destabilize the nanoparticles and hinder their tumor targeting efficiency. Additionally, the coat is functionalized with the tumor-acidity-triggered rational membrane (ATRAM) peptide for efficient and specific internalization into tumor cells in the mildly acidic microenvironment of tumors. The nanoparticles facilitated real-time fluorescence and thermal imaging of tumors and demonstrated potent NIR-light triggered anticancer activity *in vitro* and *in vivo*, without adversely affecting healthy tissue, leading to markedly prolonged survival. Our results demonstrate that the biocompatible and biodegradable ATRAM-functionalized LHAPNIRs (ALHAPNIRs) effectively combine dual-mode diagnostic imaging with targeted cancer PTT.

## INTRODUCTION

Non-invasive light-based therapies, such as photothermal therapy (PTT), have garnered considerable attention as potentially effective, fast, safe and economical alternatives to traditional cancer treatments (i.e. chemotherapy, radiotherapy and surgery).^1–3^ In cancer PTT, photothermal agents (PTAs) are used to convert absorbed light into heat, with the resulting localized hyperthermia leading to the partial or complete ablation of tumor tissue.^1,4,5^ Among the various types of PTAs currently available, fluorophores with excitation in the near-infrared (NIR) II window (950–1,350 nm) are particularly promising for PTT applications due to the greater tissue penetration, lower autofluorescence and reduced non-specific phototoxicity of NIR-II light compared to either NIR-I (700–950 nm) or visible light.^6,7^ However, clinical application of these fluorophores is often limited by their low solubility and tendency to aggregate, which attenuates the photothermal conversion efficiency, as well as their rapid *in vivo* clearance and lack of tumor specificity.^8^

A promising strategy for overcoming the limitations of current PTAs is to incorporate them into nanocarriers.^9,10^ Targeting, however, remains a major challenge for cancer drug delivery nanoplatforms, with < 1% of intravenously administered nanoparticles (NPs) ultimately reaching the target tumor tissue.^11^ A major cause of the low tumor accumulation is serum protein adsorption to the surface of NPs while in circulation.^12^ This “serum protein corona” not only destabilizes nanocarriers, it also often triggers an immune response that results in rapid blood clearance, thereby impeding tumor targeting.^13,14^ Moreover, for the small fraction of NPs that does localize to tumors, a barrier to successful delivery of their therapeutic cargo is internalization into cancer cells.^15,16^ Cellular uptake of nanocarriers occurs primarily via endocytosis, but the endosomal escape efficiency is very low (1–2%), and the majority of endocytosed NPs either undergo exocytosis or become trapped in degradative endo-lysosomal compartments.^17^

In the present study, we report development of a simple nanotheranostic system that overcomes the aforementioned issues to mediate effective tumor delivery and subsequent efficient cancer cell uptake of the photothermal fluorophore IR1061 (Figure 1). The robust nanoplatform consists of spherical hydroxyapatite NPs (HAPNs) that encapsulate the NIR dye within their pores (HAPNIRs). HAP, the principal inorganic component of human bone, possesses a number of properties that make it an attractive option as a drug delivery nanoplatform, including biocompatibility, biodegradability, tuneable pore size to modulate drug loading capacity and release kinetics, as well as ease of surface modification to increase *in vivo* circulation time and improve targeting.^18–20^ Here, the HAPNIRs are coated with a DPPC/cholesterol/DSPE-PEG bilayer, which serves to retain the encapsulated dye and allows the NPs to avoid interactions with blood serum components and evade recognition by immune cells that inhibit target recognition and result in nonspecific distribution.^21,22^ Furthermore, in order to enhance targeting, the lipid/PEG-coated HAPNIRs (LHAPNIRs) are functionalized with the acidity-triggered rational membrane (ATRAM), which facilitates specific binding to, and subsequent efficient internalization into, cancer cells in the acidic microenvironment typical of many tumors.^23–25^ Thus the ATRAM-functionalized LHAPNIRs (ALHAPNIRs) enable real-time tumor detection and monitoring via fluorescence and thermal imaging, and facilitate NIR laser light-induced PTT to substantially shrink tumors, with no noticeable systemic toxicity, thereby substantially extending survival.

**Figure 1.**
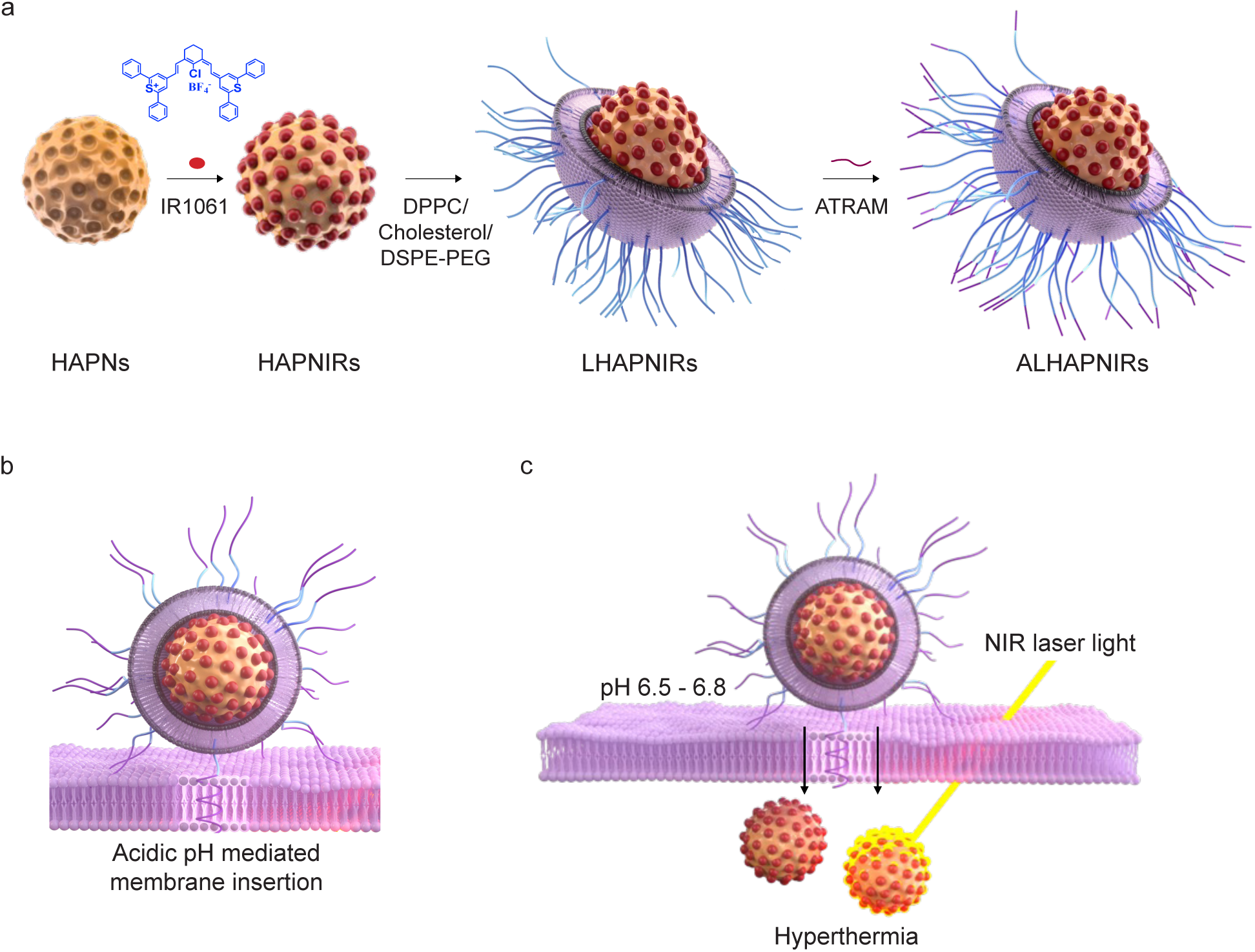
Schematic representation of preparation and mode of action of the tumor-targeted, near-infrared light-triggered, nanoplatform. (**a**) The nanoplatform consists of hydroxyapatite nanoparticles (HAPNs) that encapsulate within their pores the near-infrared (NIR) photothermal dye, IR1061. The IR1061-loaded HAPNs (HAPNIRs) are then ‘wrapped’ with a lipid/polyethylene glycol (DPPC/cholesterol/DSPE-PEG­-maleimide) coat. Finally, the lipid/PEG-coated HAPNIRs (LHAPNIRs) are functionalized with the acidity-triggered rational membrane (ATRAM) peptide (ALHAPNIRs). **(b)** In the mildly acidic conditions of the tumor microenvironment (pH ∼6.5−6.8), ATRAM promotes the targeting of ALHAPNIRs to cancer cells. (**c**) ALHAPNIRs are effectively internalized into the cancer cells, where subsequent NIR (980 nm) laser irradiation results in substantial cytotoxicity due to localized hyperthermia.

## RESULTS AND DISCUSSION

### Characterization of the synthesized hydroxyapatite nanoparticles (HAPNs) and lipid/PEG-coated HAPNs (LHAPNs)

Following synthesis (see Supporting Experimental Section), transmission electron microscopy (TEM) imaging confirmed the uniformly spherical shape of the hydroxyapatite nanoparticles (HAPNs) (Figure 2a). The X-ray diffraction (XRD) spectrum for the dried HAPNs displayed peaks consistent with previously published reports for HAP-based nanomaterials (Supporting Figure 1a).^26^ The composition of HAPNs was further verified by energy dispersive X-ray spectroscopy (EDS) and scanning transmission electron microscopy-EDS (STEM-EDS) mapping (Supporting Figures 1c and 2a). Finally, N_­_adsorption-desorption isotherms showed that HAPNs have a specific surface area of ∼23 m^2^/g and an average pore diameter of ∼1 nm (Supporting Figure 1b), which is within the range reported for other drug delivery nanoplatforms.^27^ The surface of HAPNs was coated with a DPPC/cholesterol/DSPE-PEG_­_bilayer (at a molar ratio of 77.5:20:2.5) (Figures 1a and 2b–d; Supporting Figure 2b).^25,28^ This lipid/PEG composition was selected as it was previously shown, by our group and others, to enhance colloidal stability, minimize baseline leakage, and increase *in vivo* circulation half-life.^25,28,29^ TEM and high-angle annular dark-field STEM (HAADF-STEM) imaging confirmed the presence of the lipid/PEG bilayer on the surface of HAPNs (LHAPNs) (Figure 2b–d). The uniform coating of the bilayer was further validated using TEM imaging in cryo-mode (Supporting Figure 2b). Dynamic light scattering (DLS) measurements showed that LHAPNs have a larger hydrodynamic diameter (∼89 nm) compared to uncoated HAPNs (∼78 nm) (Figure 2e; Supporting Table 1), corresponding to a bilayer coat thickness of ∼5 nm at each end of the NP, as expected. Furthermore, the zeta potential decreased from -6.5 mV to -13.2 mV following lipid/PEG coating (Figure 2f; Supporting Table 1), consistent with the values reported for other lipid-coated drug delivery NPs.^25,28^

**Figure 2.**
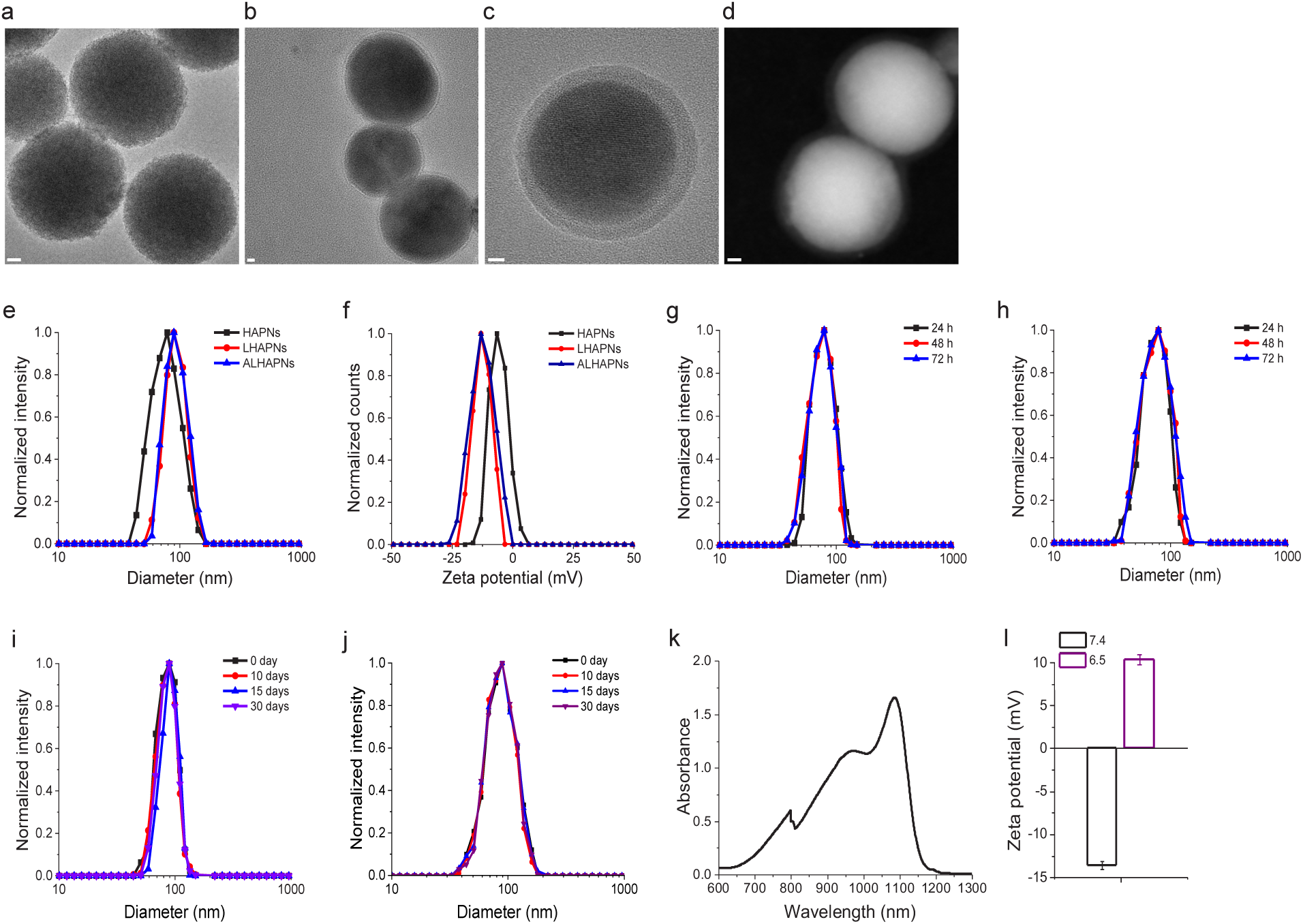
Characterization of the hydroxyapatite nanoparticles. (**a–c**) Transmission electron microscopy (TEM) images of the hydroxyapatite nanoparticles (HAPNs) (**a**) and lipid/PEG-coated HAPNs (LHAPNs) (**b**), and a magnified image of an LHAPN (**c**). (**d**) High-angle annular dark-field scanning transmission electron microscopy (HAADF-STEM) image of LHAPNs. Scale bar in **a**–**d** = 10 nm. (**e**, **f**) Size distribution analysis (**e**) and zeta potential measurements (**f**) for HAPNs, LHAPNs and ALHAPNs in 10 mM phosphate buffer (pH 7.4). (**g**, **h**) Size distribution analysis of LHAPNs in 10 mM sodium acetate buffer (pH 5.5) (**g**) and complete cell culture medium (**h**). (**i**, **j**) Long-term colloidal stability analysis of LHAPNs (**i**) and ALHAPNs (**j**) in 10 mM phosphate buffer over 30 days at 37 °C. (**k**) UV-Vis absorption spectrum of IR1061. (**l**) Comparison of the zeta potentials of ALHAPNs at pHs 7.4 and 6.5.

The colloidal stability of LHAPNs was evaluated to assess their suitability for tumor targeting and cancer therapy applications.^30,31^ The hydrodynamic diameter of the NPs remained unchanged in 10 mM phosphate buffer at pH 7.4 (89 ± 8 nm), 50 mM sodium acetate buffer at pH 5.5 (88 ± 9 nm), and complete cell culture medium (RPMI 1640, 10% fetal bovine serum (FBS), pH 7.4) (92 ± 2 nm) over 72 h (Figure 2g, h). Notably, long-term monitoring demonstrated that LHAPNs remained stable for at least a month in complete medium (∼91 nm, Figure 2i). Concomitantly, we analyzed serum protein adsorption to the surface of the NPs using quantitative proteomics (Supporting Figure 3; Supporting Table 2). After incubating HAPNs and LHAPNs in cell culture medium containing 50% FBS for 72 h, we isolated the adsorbed serum proteins and quantified them using reversed-phase liquid chromatography-tandem mass spectrometry (RPLC-MS/MS) with label-free quantification (LFQ).^32^ Analysis of the 183 most abundant serum proteins, selected after filtering the unavoidable mass spectrometric contaminants, revealed substantially higher adsorption to the HAPNs compared to the LHAPNs (Supporting Figure 3). Taken together, these studies confirm that the lipid/PEG coating enhances the colloidal stability of LHAPNs and reduces serum protein adsorption to their surface. This suggests that the designed NPs are likely to maintain an optimal size during *in vivo* circulation, thereby aiding in their targeting efficacy (i.e. tumor localization and internalization into cancer cells).^33,34^

### Photothermal properties of IR1061-Loaded LHAPs (LHAPNIRs)

Lasers operating within the second near-infrared window (NIR-II: 950–1,350 nm) offer important advantages over NIR-I (700–950 nm) and visible light lasers, including reduced absorption and scattering in biological tissues, leading to enhanced penetration and a higher energy safety threshold.^6,7^ Thus, fluorophores with excitation in the NIR-II window have gained considerable interest for tumor imaging and PTT applications.^35^ Popular amongst these PTAs is the polymethine cyanine-based organic dye IR1061, which has a maximum absorbance well within the NIR-II window and a high fluorescence quantum yield (Figure 2k).^36^ Here, we have encapsulated IR1061 within the pores of the HAPNs (HAPNIRs) using a passive entrapment loading technique (Figure 1).^37^ By adjusting the feeding ratio, a good loading capacity of IR1061 in the HAPNIRs was achieved (15 wt %; Supporting Table 3).

The photothermal properties of LHAPNIRs were investigated by measuring the temperature changes induced by NIR laser illumination of the NPs in aqueous solution. As expected, no increase in temperature was observed in the LHAPNIR samples in the absence of NIR irradiation (Figure 3a–e, g). However, upon exposure to 980 nm laser light, LHAPNIRs showed a robust photothermal response that scaled with NP concentration and irradiation power density (Figure 3a–e). For instance, at 100 µg/mL LHAPNIRs with 1.0 W/cm^2^ irradiation for 5 min the temperature rose from 28.4 ± 2.1 to 43.5 ± 1.1 °C, while at 150 µg/mL LHAPNIRs with 1.5 W/cm^2^ irradiation for 5 min the temperature increased to 55.7 ± 3.7 °C (Figure 3d, e), This suggests that the NPs loaded IR1061 can rapidly and efficiently convert NIR laser into heat of a temperature that is sufficient to ablate the cancer cells (typically ∼50 °C).^38,39^ Notably, even at low power densities,^40,41^ LHAPNIRs yielded significant temperature increases. For example, at a NP concentration of 150 µg/mL, the temperature increased to 42.1 ± 2.6 °C with 0.4 W/cm^2^ irradiation for 5 min (Figure 3e), which is comparable to the responses reported for other IR1061-loaded NPs that exhibit high photothermal conversion efficiencies.^38,39^ Moreover, the temperature increase due to NIR irradiation of LHAPNIRs was significantly greater compared to free IR1061 in saline or methanol (Figure 3f, g), which is likely due to the enhanced stability of the dye upon encapsulation in the NPs.^42^ Additionally, LHAPNIRs not only exhibited a photothermal response profile which matches that of other effective PTT nanomaterials (Figure 3i),^9,25,39,42,43^ but are also characterized by high photothermal stability, as demonstrated by the near identical maximum temperature (∼55 °C) over five successive heating/cooling cycles (Figure 3h). Together, these results underline the PTT potential of the designed NPs.

**Figure 3.**
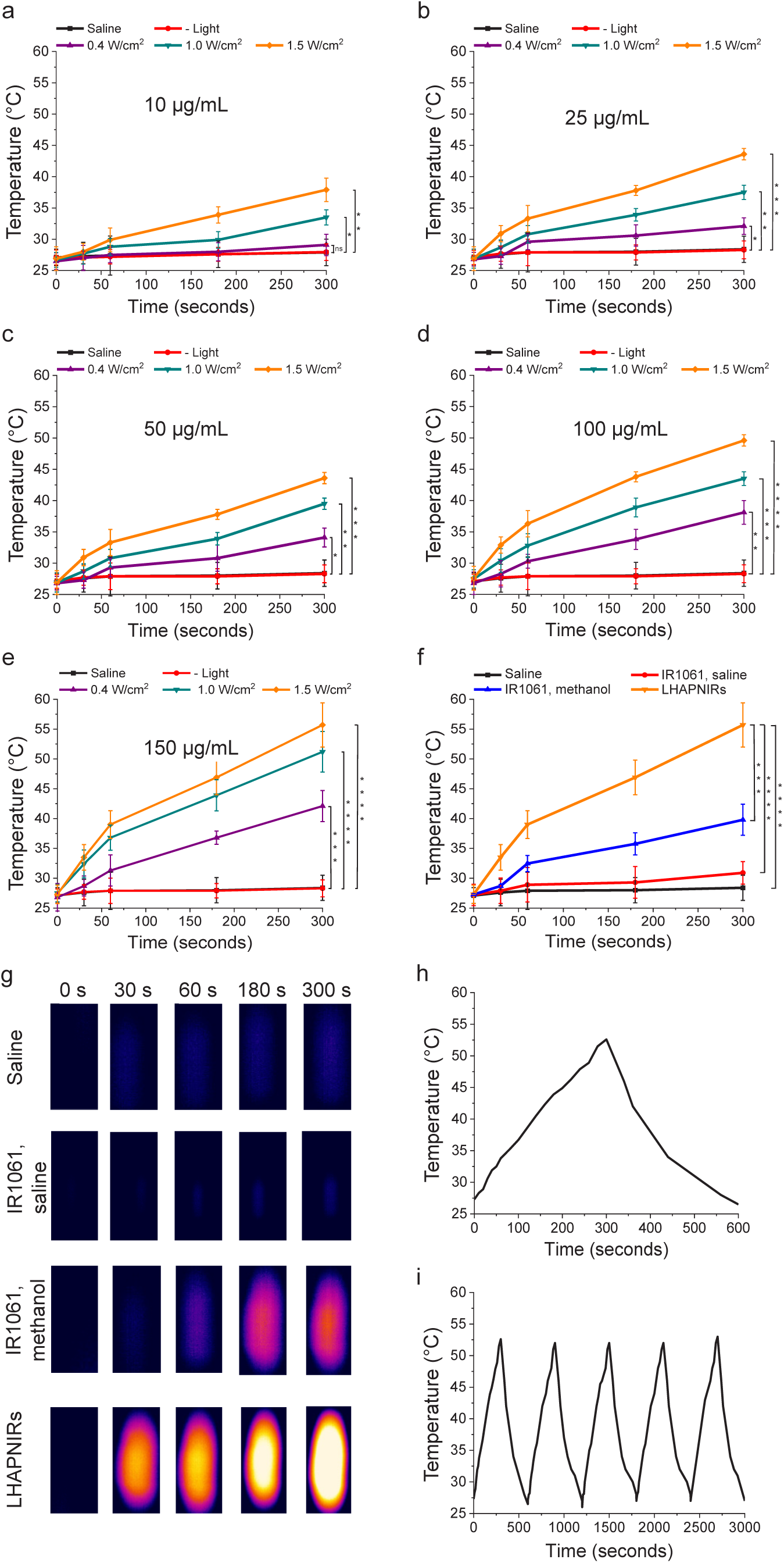
Photothermal properties of IR1061-loaded LHAPNs (LHAPNIRs). (**a**–**e**) Temperature increases following NIR laser irradiation (0.4–1.5 W/cm^2^, 5 min) of LHAPNIRs at NP concentrations of 10 µg/mL (**a**), 25 µg/mL (**b**), 50 µg/mL (**c**), 100 µg/mL (**d**) and 150 µg/mL (**e**), in 10 mM phosphate buffer (pH 7.4). (**f**) Comparison of NIR laser light (1.5 W/cm^2^, 5 min) induced temperature increases in samples of free IR1061 dye and LHAPNIRs in 10 mM phosphate buffer (pH 7.4), and free IR1061 in methanol, all measured at 0.5 µg/mL IR1061. (**g**) Thermal images of saline, IR1061 and LHAPNIRs samples illuminated with NIR laser light (1.5 W/cm^2^, 0–5 min). (**h**) Photothermal response profile of LHANIRs (150 µg/mL NPs) subjected to NIR laser irradiation (1.5 W/cm^2^, 5 min) followed by natural cooling. (**i**) Photothermal stability of LHAPNIRs (150 µg/mL NPs) monitored over 5 consecutive NIR laser irradiation (1.5 W/cm^2^, 5 min) on/off cycles. \**P* < 0.05, \*\**P* < 0.01, *\*\*\*P* < 0.001 or non-significant (ns, *P* > 0.05) for comparisons with controls or amongst the different samples.

### Cancer cell uptake of ATRAM-functionalized LHAPNs (ALHAPNs)

In order to promote specific tumor targeting, LHAPNs were functionalized with the pH-responsive acidity-triggered rational membrane (ATRAM: N_­_-CGLAGLAGLLGLEGLLGLPLGLLEGLWLGLELEGN-C_­_) peptide.^23–25,44^ This was achieved by covalently coupling the maleimide group on the PEG of the bilayer coat to the N-terminal cysteine of the peptide (see Supporting Experimental Section). At physiological pH, ATRAM remains largely unstructured and binds weakly and superficially to membranes, but in acidic conditions the peptide transitions a transmembrane α-helical conformation in lipid bilayers (Figure 1b).^23,44^ The peptide’s pH-driven membrane insertion is attributed to increased hydrophobicity resulting from protonation of its glutamate residues.^23,44^ Notably, ATRAM’s membrane insertion pKa is 6.5,^44^ making the peptide particularly well suited for targeting cancer cells in the mildly acidic solid tumor microenvironment (pH ∼6.5–6.8) (Figure 1c).^45^ As anticipated, the hydrodynamic diameter of ATRAM-functionalized LHAPNs (ALHAPNs, 90 ± 9 nm) was not appreciably larger than LHAPNs, while the zeta potential at pH 7.4 was -12.9 mV (Figure 2f; Supporting Table 1), which falls within the range reported for other highly stable nanocarriers at physiological pH.^25^ Importantly, at pH 6.5 the zeta potential of ALHAPNs becomes positive, without adversely affecting their long-term colloidal stability (Figure 2j, l; Supporting Table 1). These results strongly suggests that ALHAPNs would effectively target cancer cells within the acidic microenvironment of solid tumors.

Cellular internalization of Rhodamine B (RhB)-loaded ALHAPNs (ALHAPNRhBs) in MIA PaCa-2 cancer cells was assessed using confocal fluorescence microscopy and flow cytometry (Figure 4a–d). Human pancreatic MIA PaCa-2 cells were treated with ALHAPNRhBs for 4 h at pH 7.4 or 6.5. Confocal microscopy imaging clearly showed substantially higher cellular uptake and cytosolic localization of the NPs under acidic conditions relative to physiological pH (Figure 4a). Consistent with the imaging results, flow cytometry analysis revealed that internalization of ALHAPNRhBs at pH 6.5 was ∼9-fold higher than that at pH 7.4 (Figure 4b–d). In marked contrast, the internalization of LHAPNRhBs (i.e. in the absence of ATRAM) was poor at both pHs (7.4 and 6.5) (Figure 4b–d). These results demonstrate that ATRAM effectively drives uptake of the NPs specifically in cancer cells within a mildly acidic environment.

**Figure 4.**
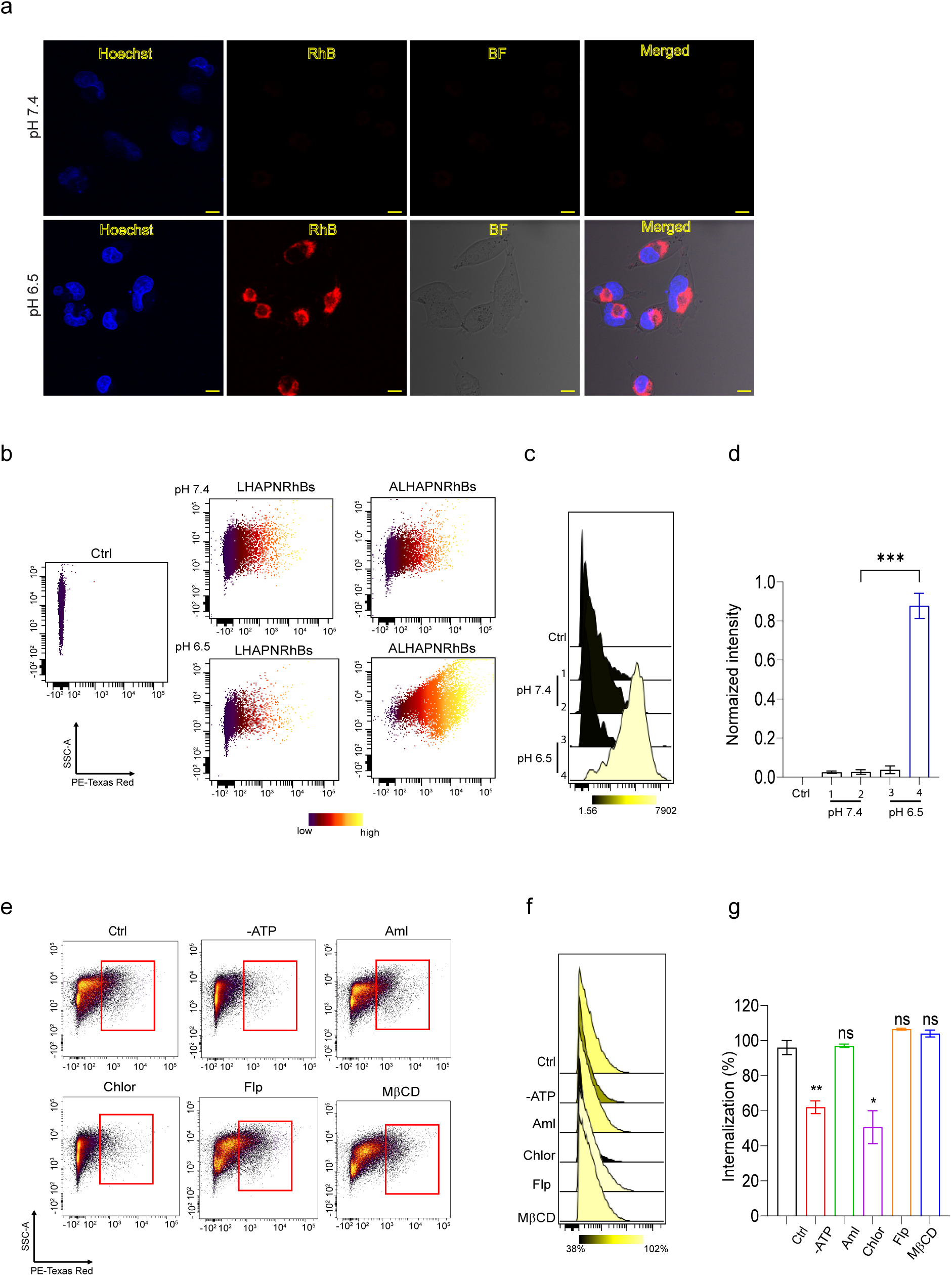
pH-dependent cellular uptake of dye-loaded ALHAPNs. **(a)** Confocal fluorescence microscopy images of MIA PaCa-2 cells incubated with Rhodamine B (RhB)-loaded ALHAPNs (ALHAPNRhBs) (0.5 µg/mL RhB) for 4 h at physiological (*top panels*) or acidic (*lower panels*) pH. RhB is red-coloured for clarity. Imaging experiments were performed in triplicate and representative images are shown. Scale bar = 10 µm. **(b–d)** Flow cytometry analysis of MIA PaCa-2 cells treated with the RhB-loaded LHAPNs without or with ATRAM (LHAPNRhBs or ALHAPNRhBs, respectively) at pH 7.4 or 6.5. Shown are the flow cytometry dot plots (**b**) and histograms (**c**), as well as bar graph summary (*n* = 3) of the uptake of LHAPNRhBs (bar 1, pH 7.4; bar 3, pH 6.5) and ALHAPNRhBs (bar 2, pH 7.4; bar 4, pH 6.5) (**d**). **(e–g)** Cellular uptake mechanisms of ALHAPNRhBs. Flow cytometry dot plots (**e**), histograms (**f**), and bar graph summary (*n* = 3) (**g**), of cellular uptake of the NPs in MIA PaCa-2 cells pretreated with sodium azide/2-deoxy-D-glucose to deplete cellular ATP (-ATP), or treated with endocytosis inhibitors – amiloride (Aml; macropinocytosis inhibitor), chlorpromazine (Chlor; clathrin-mediated endocytosis), filipin (Flp; caveolae-dependent endocytosis), or methyl-β-cyclodextrin (MβCD; lipid raft-associated endocytosis) – compared with uninhibited uptake in control cells (Ctrl). \**P* < 0.05, \*\**P* < 0.01, *\*\*\*P* < 0.001 or non-significant (ns, *P* > 0.05) for comparisons with controls or amongst the different samples.

Next the cellular internalization mechanism(s) of ALHAPNRhBs was determined using a series of flow cytometry experiments at pH 6.5 (Figure 4e–g). First, intracellular ATP was depleted with sodium azide/deoxyglucose, which resulted in a partial reduction in the uptake of ALHAPNRhBs. This suggests that the NPs are internalized through both energy-dependent (endocytosis) and independent (direct translocation) mechanisms (Figure 4e−g). Direct translocation likely involves ATRAM-driven membrane anchoring, followed by the fusion of the lipid/PEG coat with the cancer cell membrane, resulting in the release of the NPs into the cytosol.^24^ To further elucidate the nature of the energy-dependent pathway, we pretreated cells with specific inhibitors of endocytosis: amiloride (which blocks micropinocytosis by inhibiting Na^+^/H^+^ exchange, chlorpromazine (which inhibits clathrin-coated pit formation), filipin (an inhibitor of caveolae-dependent endocytosis), and methyl-β-cyclodextrin (which disrupts lipid raft-associated endocytic pathways by depleting cholesterol from the plasma membrane).^46–49^ Among the endocytosis inhibitors tested, only chlorpromazine significantly reduced the cellular uptake of ALHAPNRhBs, indicating that clathrin-mediated endocytosis (CME) contributes to the NP’s cellular internalization (Figure 4e−g).

In the case of both cellular internalization pathways, ALHAPNs gain access to the cytosol of cancer cells: direct translocation across the plasma membrane provides ALHAPNs with direct access to the cytosol; likewise, after CME, the acidification of mature endocytic compartments facilitates endosome membrane insertion and disruption by ATRAM, akin to the behaviour of other pH-responsive peptides,^25,50,51^ resulting in cytosolic release of the NPs. Thus, the multi-mode uptake at acidic pH allows ALHAPNs to efficiently enter tumor cells.

### Cancer cell toxicity of IR1061-loaded ALHAPNs (ALHAPNIRs)

The anticancer potency of the designed NPs was assessed using the MTS cell viability assay.^52,53^ Treatment with IR1061-free LHAPNs (5–150 µg/mL) did not reduce MIA PaCa-2 cell viability in the absence or presence of NIR laser irradiation (Supporting Figure 4a). Likewise, without exposure to NIR laser light, LHAPNIRs were not toxic to MIA PaCa-2 cells, up to an IR1061 concentration of 5 µg/mL, either at physiological or acidic pH (Supporting Figure 4b). These results confirm the biocompatibility of the NPs and, in turn, their suitability for cancer imaging and PTT applications.

In the presence of 980 nm laser light, treatment with ALHAPNIRs for 24–72 h at pH 7.4 had no adverse effect on the viability of MIA PaCa-2 cells (Figure 5a–f and Supporting Figures 5 and 6), which can be attributed to the poor cell internalization of the NPs at physiological pH (Figure 4a, b). Conversely, treatment with ALHAPNIRs for the same duration at pH 6.5 substantially reduced viability of the cells, and the cytotoxicity of the NPs scaled with IR1061 concentration and laser power density/irradiation duration (Figure 5a–f and Supporting Figures 5 and 6). Consistent with the MTS assay results, calcein AM/propidium iodide (PI) staining (an established method for visually detecting live/dead cells)^54,55^ of MIA PaCa-2 cells showed that combining ALHAPNIR treatment at pH 6.5 and 980 nm laser irradiation yielded a clear decrease in the fraction of live cells (calcein AM signal), with a concomitant increase in the fraction of dead cells (PI signal) (Figure 6a).

**Figure 5.**
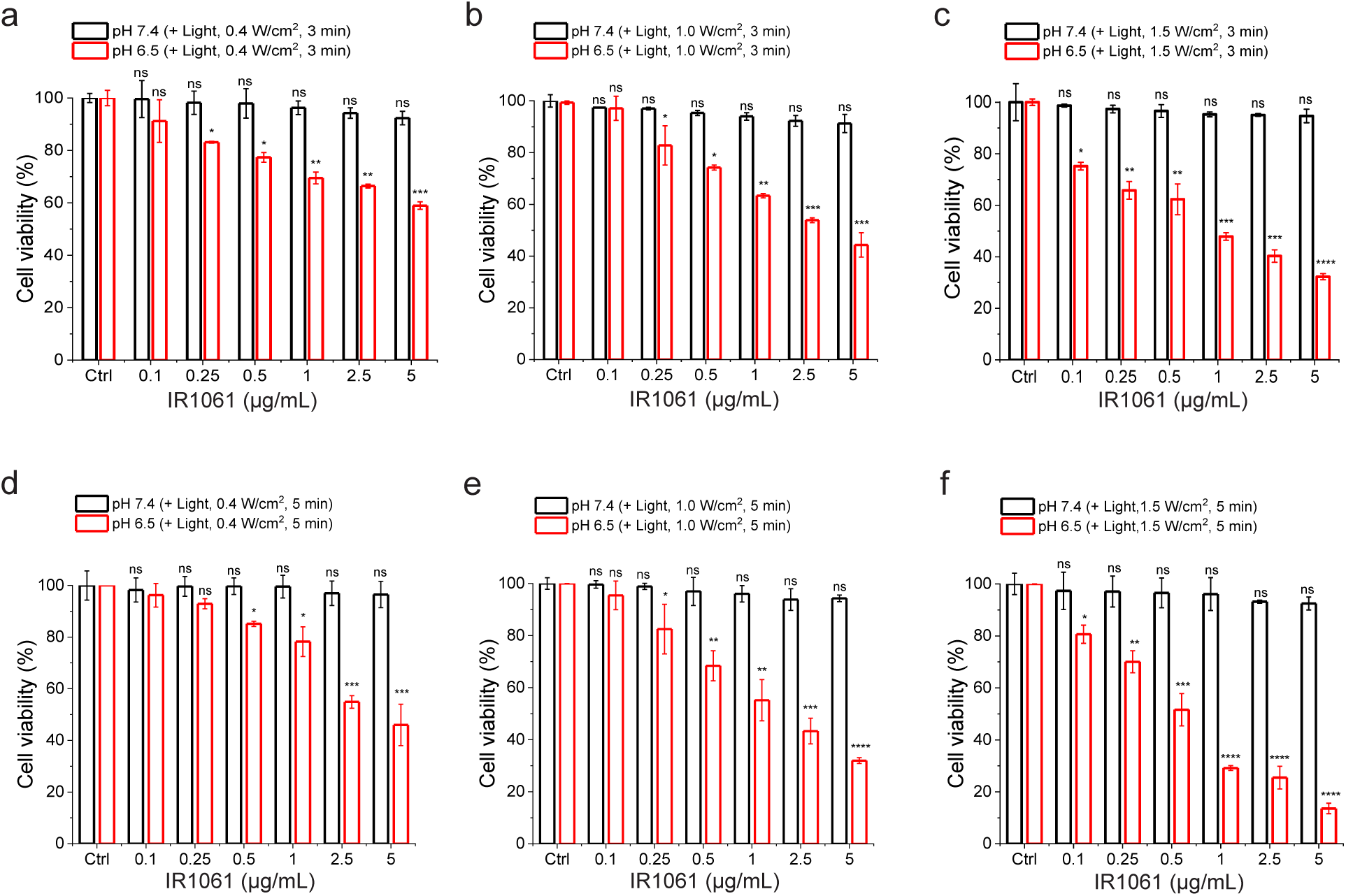
NIR-light triggered cytotoxicity of ALHAPNIRs. Cell viability of MIA PaCa-2 cells treated with ALHAPNIRs (0.1–5 µg/mL IR1061) for 48 h and exposed to NIR laser light of different irradiation power densities (0.4–1.5 W/cm^2^) for 3.0 (**a–c**) or 5.0 (**d–f**) min at pH 7.4 or 6.5. Cell viability in was measured using the MTS assay, with the % viability determined form the ratio of the absorbance of the treated cells to the control cells (*n* = 3). *\*P* < 0.05*, **P* < 0.01*, ***P* < 0.001*, ****P* < 0.0001 or non-significant (ns, *P* > 0.05) compared with controls.

**Figure 6.**
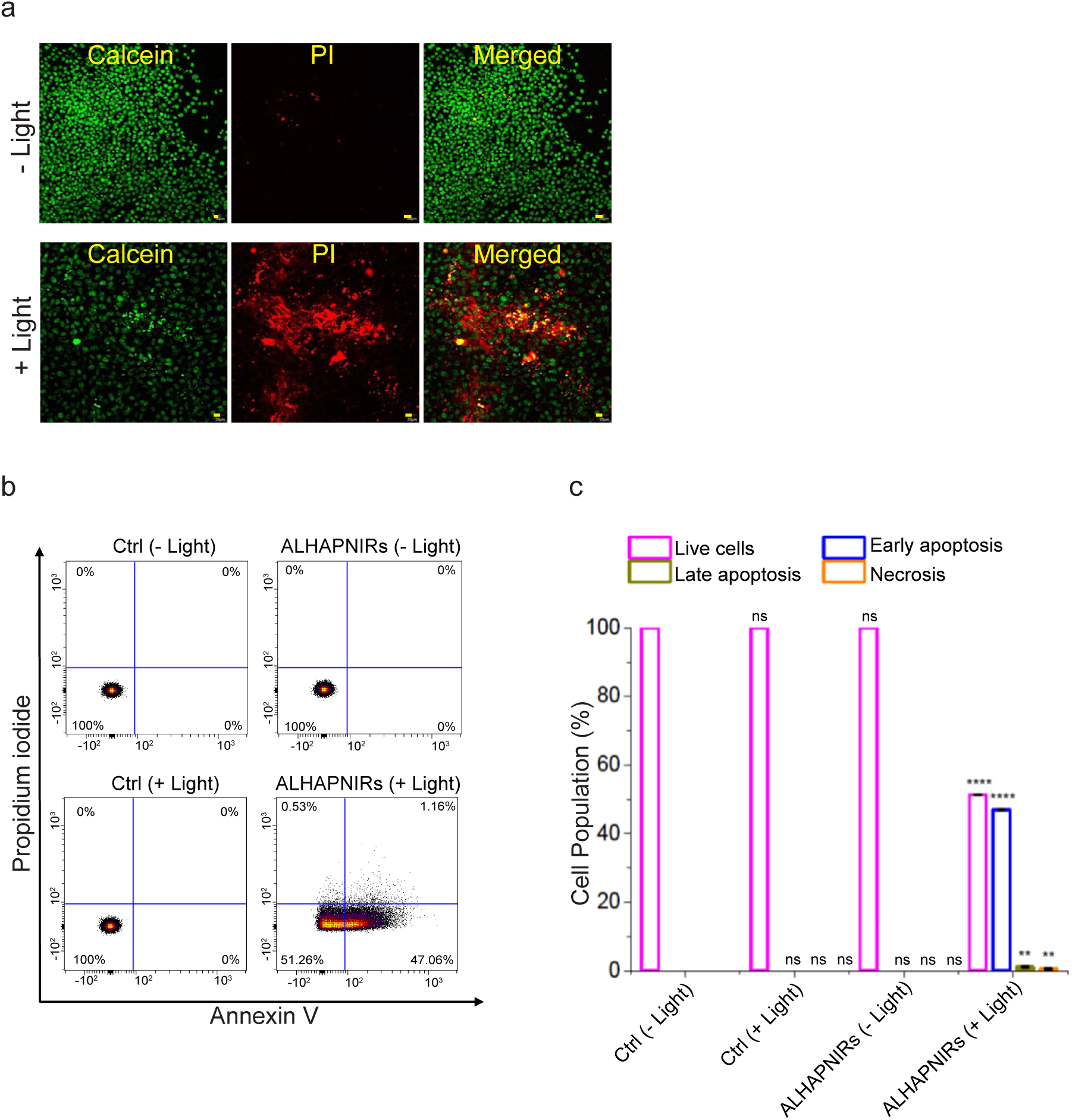
Mechanism of ALHAPNIR-induced cytotoxicity. (**a**) Confocal fluorescence microscopy images of calcein AM/ propidium iodide (PI) staining of MIA PaCa-2 cells incubated with ALHAPNIRs (0.5 µg/mL IR1061) for 12 h at pH 6.5 in the absence (-Light) or presence (+ Light) of NIR laser irradiation (1.0 W/cm^2^, 5 min). Scale bar = 20 μm. The experiments were performed in triplicate and representative images are shown. (**b, c**) Flow cytometry analysis of annexin V/PI-stained MIA PaCa-2 cells that were either untreated (control, Ctrl), or treated with ALHAPNIRs (0.5 µg/mL IR1061) for 48 h at pH 6.5, with or without exposure to NIR light (1.5 W/cm^2^, 5 min). The four quadrants are defined as follows: annexin V^-^/PI^-^ (bottom left), live cells; annexin V^+^/PI^-^ (bottom right), early apoptotic cells; annexin V^+^/PI^+^ (top right), late apoptotic cells; and annexin V^-^/PI^+^ (top left), necrotic cells. (**c**) A summary of the incidence of early/late apoptosis and necrosis in the MIA PaCa-2 cells treated with ALHAPNIRs determined from the flow cytometry analysis of annexin V/PI staining in (**b**) (*n* = 3). *\*P* < 0.05*, **P* < 0.01*, ***P* < 0.001*, ****P* < 0.0001 or non-significant (ns, *P* > 0.05) compared with controls.

Finally, staining of MIA PaCa-2 cells with FITC-conjugated annexin V/PI followed by sorting with flow cytometry^24,25,56–58^ revealed that exposure to ALHAPNIRs (0.5 µg/mL IR1061) and 980 nm laser irradiation (1.0 W/cm^2^, 5 min) at pH 6.5 led to a sizeable fraction of the cells undergoing apoptosis (Figure 6b, c). Of note, the viability obtained by annexin V/PI staining under these conditions was 51.3 ± 0.01%, which is in excellent agreement with the value measured using the MTS assay under the same conditions (51.7 ± 6.2%) (Figures 5f and 6c). Taken together, the cell uptake and viability/toxicity experiments demonstrate that ALHAPNIRs cause extensive NIR light-triggered toxicity, due to hyperthermia-induced apoptosis, selectively in malignant cells within a mildly acidic environment (Figures 3–6).

### Macrophage recognition and immunogenicity of ALHAPNIRs

Opsonization and subsequent uptake by monocytes and macrophages of the mononuclear phagocyte system (MPS) results in the accumulation of NPs in healthy organs, like the spleen and liver, rather than in target tumors.^59^ To address this challenge, NPs are often coated with neutral molecules, such as PEG, which are known to reduce serum protein adsorption and clearance by the MPS.^21^

To ascertain the potential of the lipid/PEG coat to prevent premature clearance the IR1061-loaded HAPNs by the MPS, we assessed the interaction of the NPs with differentiated human monocytic leukemia THP-1 cells, a widely used model of monocyte/macrophage activation.^60^ Treatment of differentiated THP-1 cells with HAPNIRs lead to a significant decrease in viability and markedly elevated production of the inflammatory cytokines tumor necrosis factor-alpha (TNF-α) and interlukin 1 beta (Il-1β) (Supporting Figure 7a, b). However, negligible toxicity and TNF-α/Il-1β production was detected following exposure of the cells to ALHAPNIRs under the same experimental conditions (Supporting Figure 7a, b). These results demonstrate that the lipid/PEG coat not only prevents the formation of a serum protein corona on the surface of the NPs (Supporting Figure 3), it also minimizes interactions with immune cells (Supporting Figure 7), properties which are critical for effective tumor targeting.

### Tumor localization of dye-loaded ALHAPNs

MIA PaCa-2 tumor-bearing mice were intravenously injected with NPs encapsulating the far-red lipophilic carbocyanine DiD (HAPNDiDs, LHAPNDiDs or ALHAPNDiDs) and subsequently imaged using the IVIS Spectrum optical and thermal imaging system at various time-points (0–72 h) post injection. Fluorescence imaging showed substantially higher accumulation of ALHAPNDiDs in tumors compared to LHAPNDiDs or HAPNDiDs (Figure 7a, b). The peak tumor localization of ALHAPNDiDs occurred at 8 h, but the NPs persisted in tumor tissue up to 72 h post *i.v.* injection (Figure 7a, b). This was confirmed by *ex vivo* fluorescence imaging of MIA PaCa-2 tumors excised at 72 h post injection (Figure 7c). Importantly, *ex vivo* imaging also revealed negligible presence of ALHAPNDiDs in the vital organs (lungs, liver, spleen, kidney or heart) (Figure 7c).

**Figure 7.**
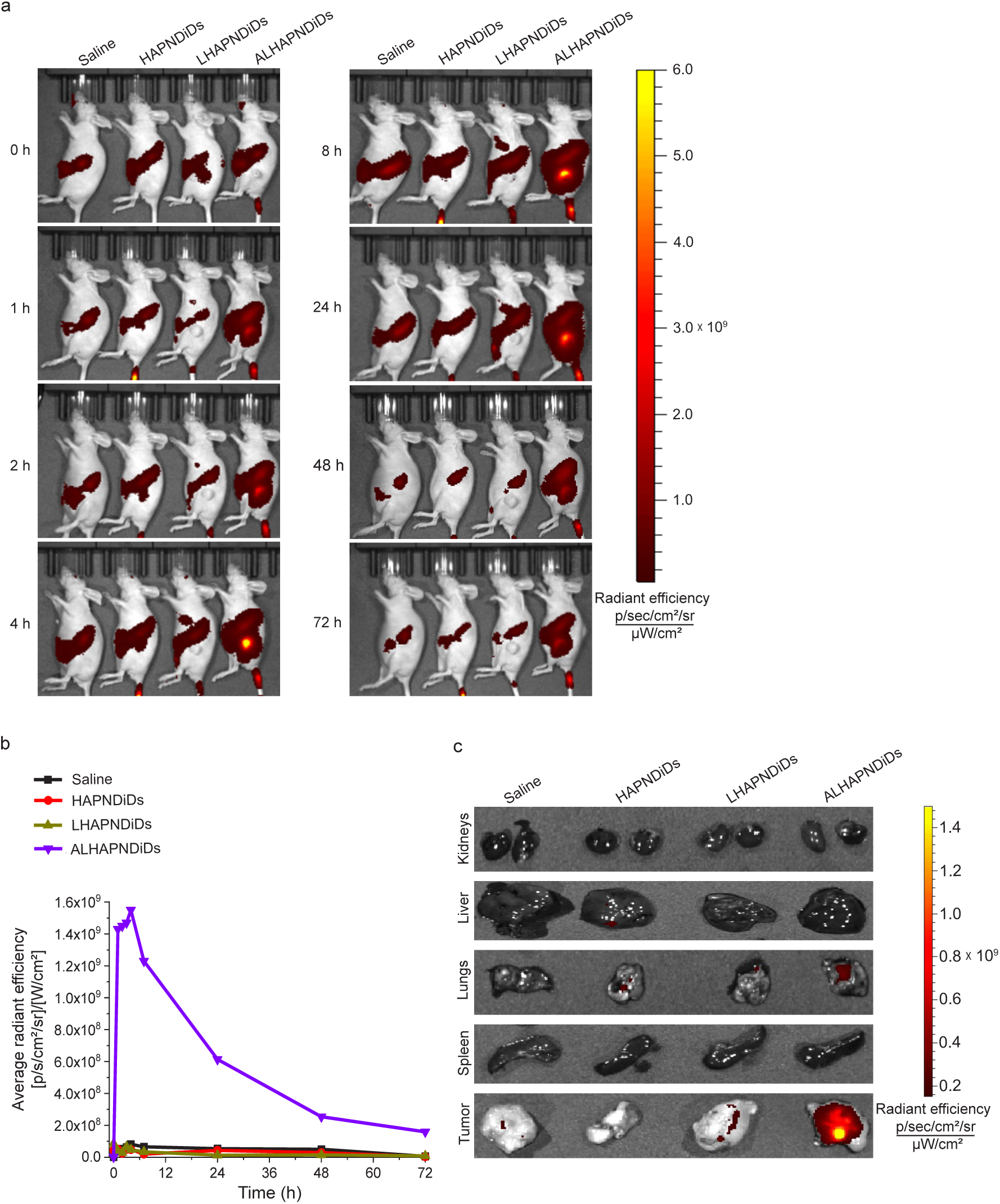
*In vivo* tumor localization of dye-loaded ALHAPNs. (**a**) *In vivo* biodistribution of NPs encapsulating the far-red lipophilic fluorophore DiD (HAPNDiDs, LHAPNDiDs and ALHAPNDiDs; 5 mg/kg NPs, 0.3 mg/kg DiD)^66^ in MIA PaCa-2 tumor bearing mice. The images were acquired at different time points (0–72 h) following *i.v.* injection of the NPs. (**b**) *In vivo* quantification of fluorescence intensity (expressed in terms of average radiant efficiency) in tumor tissue. Data was acquired by whole-mouse imaging (*n* = 4 per group). (**c**) *Ex vivo* imaging of tumors and vital organs (kidneys, liver, lungs, and spleen) isolated from mice 72 h after *i.v.* injection of HAPNDiDs, LHAPNDiDs, and ALHAPNDiDs.

Consistent with the fluorescence imaging results, thermal imaging of MIA PaCa-2 tumor-bearing mice following 980 nm laser irradiation (1.0 W/cm^2^) at 8 h post *i.v.* injection showed that ALHAPNIRs induced a much more rapid and pronounced temperature increase in the tumors (from 36 to 45 °C within 5 min) compared to HAPNIRs and LHAPNIRs (Supporting Figure 8). Indeed, this photothermal response is comparable to that of other highly effective PTT nanomaterials,^61,62^ which underscores the high *in vivo* photothermal conversion efficiency and photostability of ALHAPNIRs. These results illustrate that ALHAPNIRs effectively target tumors and facilitate fluorescence and thermal imaging of the cancerous tissue.

### *In vivo* anticancer effects of ALHAPNIRs

MIA PaCa-2 tumor-bearing mice were administered intravenous injections of HAPNIRs, LHAPNIRs or ALHAPNIRs (22 mg/kg NPs, 3.3 mg/kg IR1061) once every 2 days for a total of 15 doses (Figure 8a, b). The dosage of IR1061 injected was comparable to that used in other PTT-based cancer treatment studies.^43^ As anticipated, in the absence of 980 nm laser irradiation, none of the treatments significantly affected tumor growth or overall survival of the mice (Figure 8d, f, g, h). In the presence of NIR laser irradiation, HAPNIR treatment had a negligible effect on tumor growth and survival (Figure 8e, f, g, i). More pronounced effects, in terms of reduced tumor growth and increased median survival time, were observed in the LHAPNIR treatment group (Figure 8e, f, g, i). Notably, treatment with ALHAPNIRs yielded the greatest anticancer effects, completely reversing tumor growth and markedly prolonging survival compared to the controls and the other treatment groups over the duration of the experiment (Figure 8e, f, g, i).

**Figure 8.**
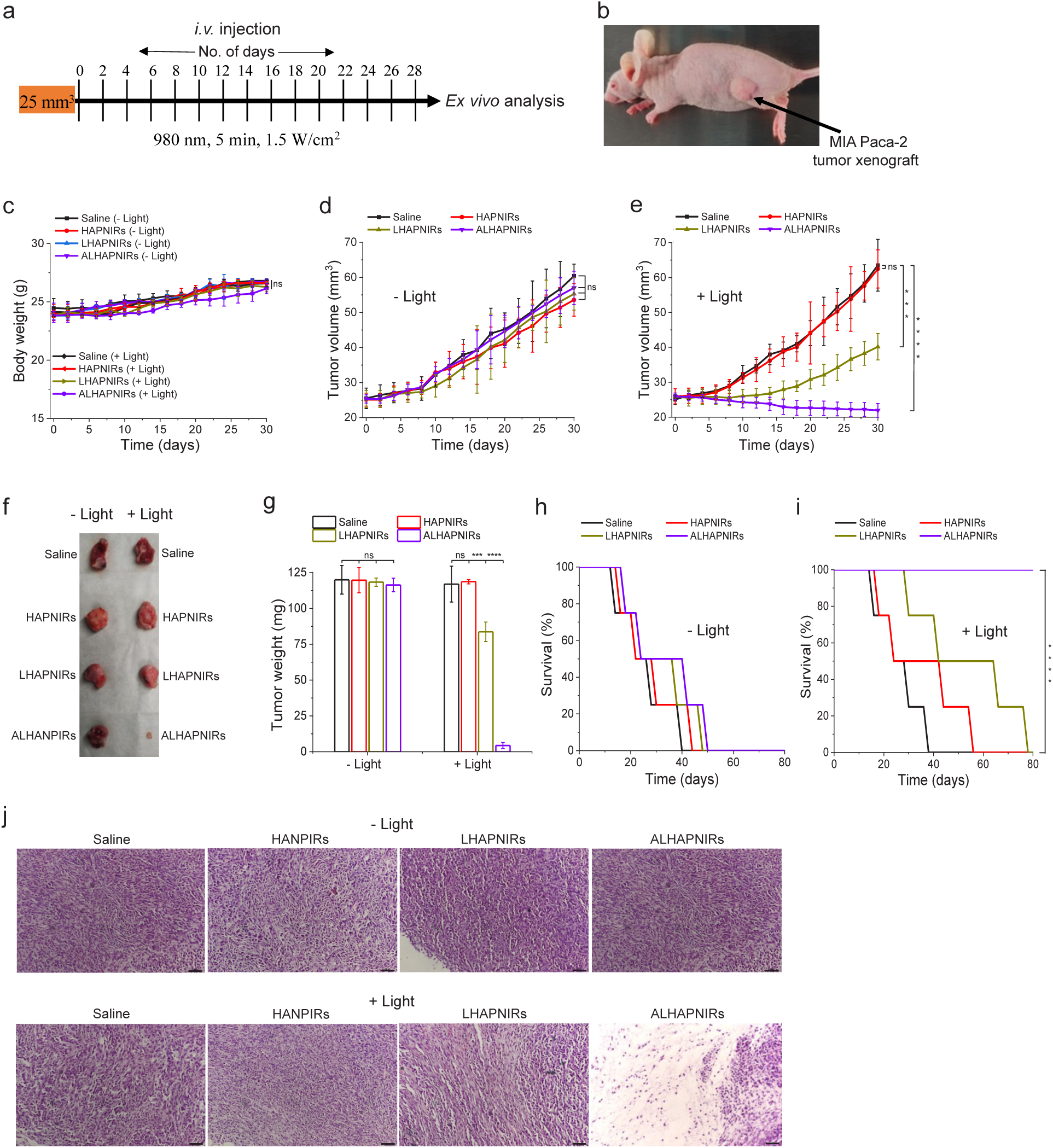
Inhibition of tumor growth by ALHAPNIRs. (**a**) Design of the tumor reduction studies. Once the MIA PaCa-2 tumor volume reached *∼*25 mm^3^, the mice were randomized into the different treatment groups (*n* = 16 per group), which were injected intravenously with saline, HAPNIRs, LHAPNIRs or ALHAPNIRs (22.5 mg/kg NPs, 3.3 mg/kg IR1061). Injections were done every other day for a total of 15 doses, with the first day marked at day 0. Within each treatment group, 8 mice were subjected to NIR laser irradiation (980 nm, 1.5 W/cm^2^, 5 min) at 8 h post injection. (**b**) Image of MIA PaCa-2 tumor-bearing mouse at day 0. (**c**) Body weight changes of the MIA PaCa-2 tumor-bearing mice in the different treatment groups in the absence (- Light) or presence (+ Light) of irradiation monitored for the duration of the experiment. (**d, e**) Tumor volume measurements for the saline, HAPNIR, LHAPNIR and ALHAPNIR groups over 30 days of treatment in the absence (- Light) or presence (+ Light) of NIR light irradiation (*n* = 8 per group). (**f, g**) Tumor mass measurements for the different treatment groups. After 30 days of treatment, 4 mice per treatment group were sacrificed, and the tumor tissues were isolated and imaged (**f**) and weighed to determine (**g**). (**h, i**) Survival curves for the different treatment groups (saline, HAPNIRs, LHAPNIRs and ALHAPNIRs) over 80 days in the absence (**h**) or presence (**i**) of NIR light irradiation (*n* = 4 per group). (**j**) H&E staining of tumor sections from the different groups after the 30 days of treatment in the absence (top panels) or presence (lower panels) of NIR light irradiation. Images shown in (**j**) are representative of tissue sections from 4 mice per treatment group; scale bar = 50 μm. *\*\*\*P* < 0.001*, ****P* < 0.0001 or non-significant (ns, *P* > 0.05) compared with controls.

Histological analysis using hematoxylin and eosin (H&E) staining corroborated the superior antitumor efficacy of ALHAPNIRs relative to other treatments (Figure 8j). Importantly, treatment with ALHAPNIRs, either with or without NIR irradiation, did not adversely affect the body weight of the mice (Figure 8c), and no abnormalities or lesions were detected in H&E-stained sections of vital organs (heart, kidneys, liver, lungs, and spleen) following treatment (Supporting Figure 9). Additionally, exposure to ALHAPNIRs did not significantly increase the concentrations of the inflammatory cytokines TNF-α and Il-1β in circulation, which confirms the lack of systemic toxicity of the NPs (Supporting Figure 10). Collectively, these findings compellingly demonstrate that IR1061-loaded ALHAPNs effectively target and shrink tumors, through NIR-induced PTT, while safeguarding healthy tissue and significantly enhancing survival outcomes.

## CONCLUSIONS

PTT holds great promise as a non-invasive cancer treatment modality.^1,2,43,63^ However, several challenges have hindered its clinical application. Key issues include poor solubility, low circulation stability, lack of target specificity, and inefficient cellular uptake of PTAs. Here, we report a simple and efficacious nanotheranostic solution: ATRAM-functionalized, lipid/PEG-coated HAP nanoparticles encapsulating the NIR-II photothermal dye IR1061 (ALHAPNIRs). The biocompatible and biodegradable ALHAPNIRs overcome the limitations of many current PTAs by combining the following critical properties: (i) stable encapsulation of PTAs, which effectively inhibits their aggregation and prevents their premature degradation, allowing for sustained therapeutic activity over an extended period; (ii) minimal interactions with healthy tissue, serum proteins and macrophages, thereby appreciably increasing the *in vivo* circulation time and, in turn, tumor accumulation of the PTA cargo; (iii) efficient and specific uptake into cancer cells within the mildly acidic microenvironment of solid tumors, which is crucial for maximizing therapeutic efficacy while minimizing damage to surrounding healthy tissue; (iv) potent localized PTT triggered by NIR-II light, which has greater tissue penetration, lower autofluorescence and reduced phototoxicity compared to either NIR-I or visible light; (v) NIR-mediated dual-mode – fluorescence and thermal – imaging, enabling simultaneous localization of lesions and real-time monitoring of therapeutic effects, thereby enhancing both diagnostic accuracy and therapeutic precision.^64,65^ Collectively, these features underscore the potential of ALHAPNIRs as a promising nanoplatform that offers robust diagnostic imaging and targeted PTT capabilities in a single, integrated system.

## ACKNOWLEDGEMENTS

The authors thank Ms. Jumaanah Alhashemi, Assistant Director of Research Visualization and Fabrication Services at NYU Abu Dhabi, for creating the graphic illustrations, and Mohayed Magzoub (MEng MSc MA CEng MIMarEST RN) for critical reading of the manuscript. The authors also acknowledge the NYU Abu Dhabi Center for Genomics and Systems Biology (NYUAD-CGSB) for granting access to the BD FACSAria III for flow cytometry measurements. Imaging – confocal fluorescence microscopy, TEM, and IVIS Spectrum – and Zetasizer measurements, were conducted using the Core Technology Platforms (CTP) at NYU Abu Dhabi. Additionally, proteomics data processing utilized the High-Performance Computing (HPC) resources at NYU Abu Dhabi. This research was funding from NYU Abu Dhabi (grant AD389) to M.M., and from the U.S. National Institutes of Health (R35GM140846) to F.N.B.

## SUPPORTING INFORMATION

**Supporting Figures:** characterization of the hydroxyapatite nanoparticles (HAPNs); electron microscopy imaging of HAPNs and lipid/PEG-coated HAPNs (LHAPNs); quantitative proteomic analysis of serum protein adsorption to the surface of HAPNs and LHAPNs; cell viability control experiments; NIR-light triggered cytotoxicity of the ATRAM-functionalized LHAPNIRs (ALHAPNIRs) at 24 h incubation; NIR-light triggered cytotoxicity of ALHAPNIRs at 72 h incubation; macrophage recognition and immunogenicity of the NPs; ALHAPNIR-facilitated photothermal tumor imaging; histological analysis of vital organs following treatment with the NPs; quantification of inflammatory cytokines in circulation following treatment with the NPs.

**Supporting Tables:** summary of hydrodynamic diameters and zeta potentials of the hydroxyapatite (HAP) NPs; proteins corresponding to the UniProt Knowledgebase (UniProtKB) accession numbers shown in Supporting Figure 3; IR1061 dye loading capacity of the hydroxyapatite nanoparticles (HAPNs).

**Supporting Experimental Section:** materials; synthesis of the hydroxyapatite nanoparticles; characterization of the nanoparticles; quantitative proteomic analysis; photothermal response; cell culture; cancer cell uptake; cell viability/toxicity assays; macrophage toxicity and immunogenicity; *in vivo* experiments; biodistribution and tumor accumulation; tumor growth inhibition; statistical analysis.

