## Supporting Information for "Tumor-targeted hydroxyapatite nanoparticles for dual-mode diagnostic imaging and near-infrared light-triggered photothermal cancer therapy"

### TABLE OF CONTENTS

#### SUPPORTING FIGURES


Supporting Figure 1. Characterization of the hydroxyapatite nanoparticles (HAPNs)

Supporting Figure 2. Electron microscopy imaging of HAPNs and lipid/PEG-coated HAPNs (LHAPNs)

Supporting Figure 3. Quantitative proteomic analysis of serum protein adsorption to the surface of HAPNs and LHAPNs

Supporting Figure 4. Cell viability control experiments

Supporting Figure 5. NIR-light triggered cytotoxicity of the ATRAM-functionalized LHAPNIRs (ALHAPNIRs) at 24 h incubation

Supporting Figure 6. NIR-light triggered cytotoxicity of ALHAPNIRs at 72 h incubation

Supporting Figure 7. Macrophage recognition and immunogenicity of the NPs

Supporting Figure 8. ALHAPNIR-facilitated photothermal tumor imaging

Supporting Figure 9. Histological analysis of vital organs following treatment with the NPs

Supporting Figure 10. Quantification of inflammatory cytokines in circulation following treatment with the NPs


#### SUPPORTING TABLES


Supporting Table 1. Summary of hydrodynamic diameters and zeta potentials of the hydroxyapatite (HAP) NPs

Supporting Table 2. Proteins corresponding to the UniProt Knowledgebase (UniProtKB) accession numbers shown in Supporting Figure 3

Supporting Table 3. IR1061 dye loading capacity of the hydroxyapatite nanoparticles (HAPNs)

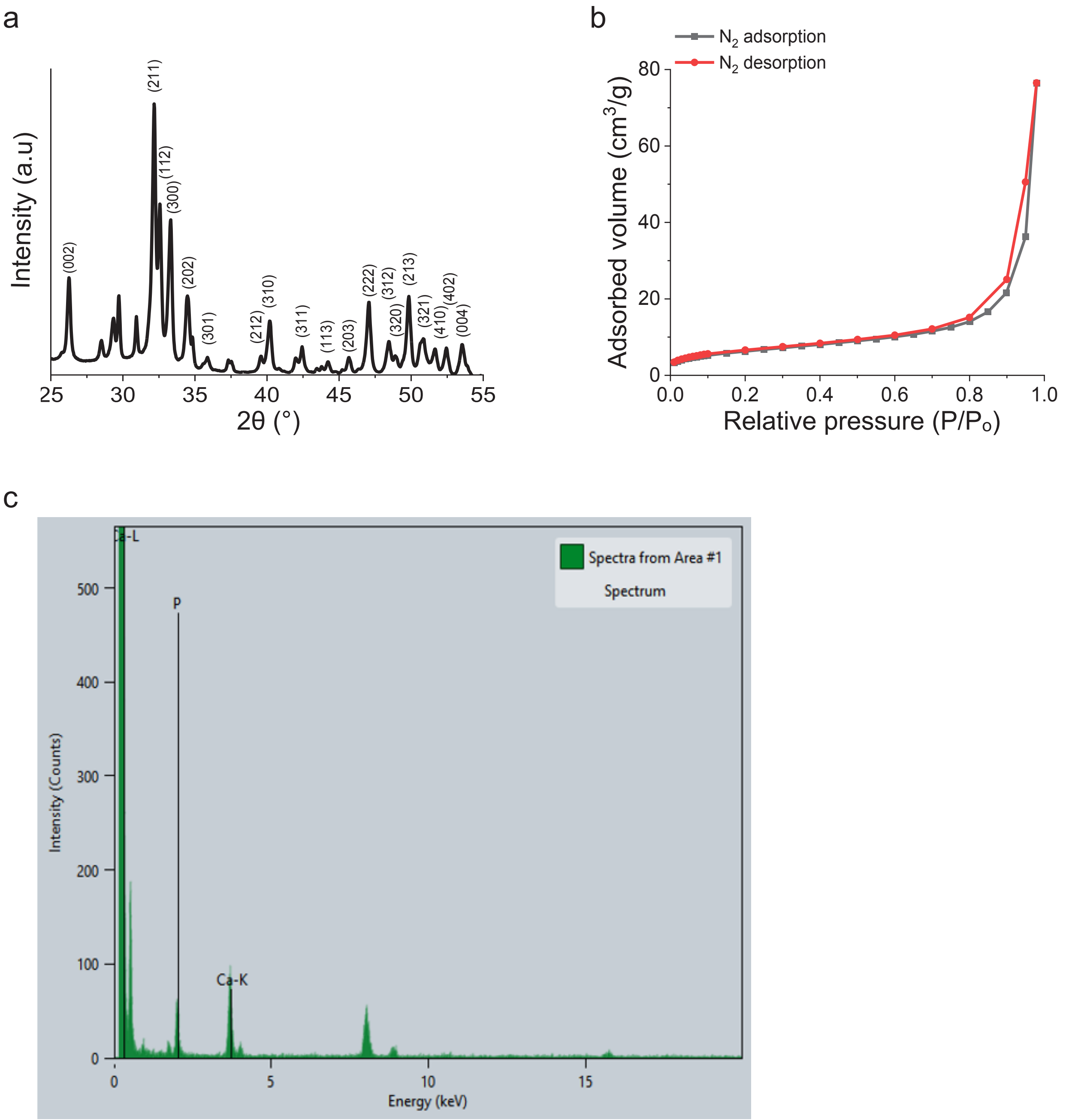


**Supporting Figure 1. Characterization of the hydroxyapatite nanoparticles (HAPNs).** (**a**) Powder X-ray diffraction (XRD) spectrum of the HAPNs. (**b**) $N_2$ adsorption-desorption isotherms of the HAPNs. (**c**) Energy dispersive X-ray spectroscopy (EDS) spectrum of the HAPNs.

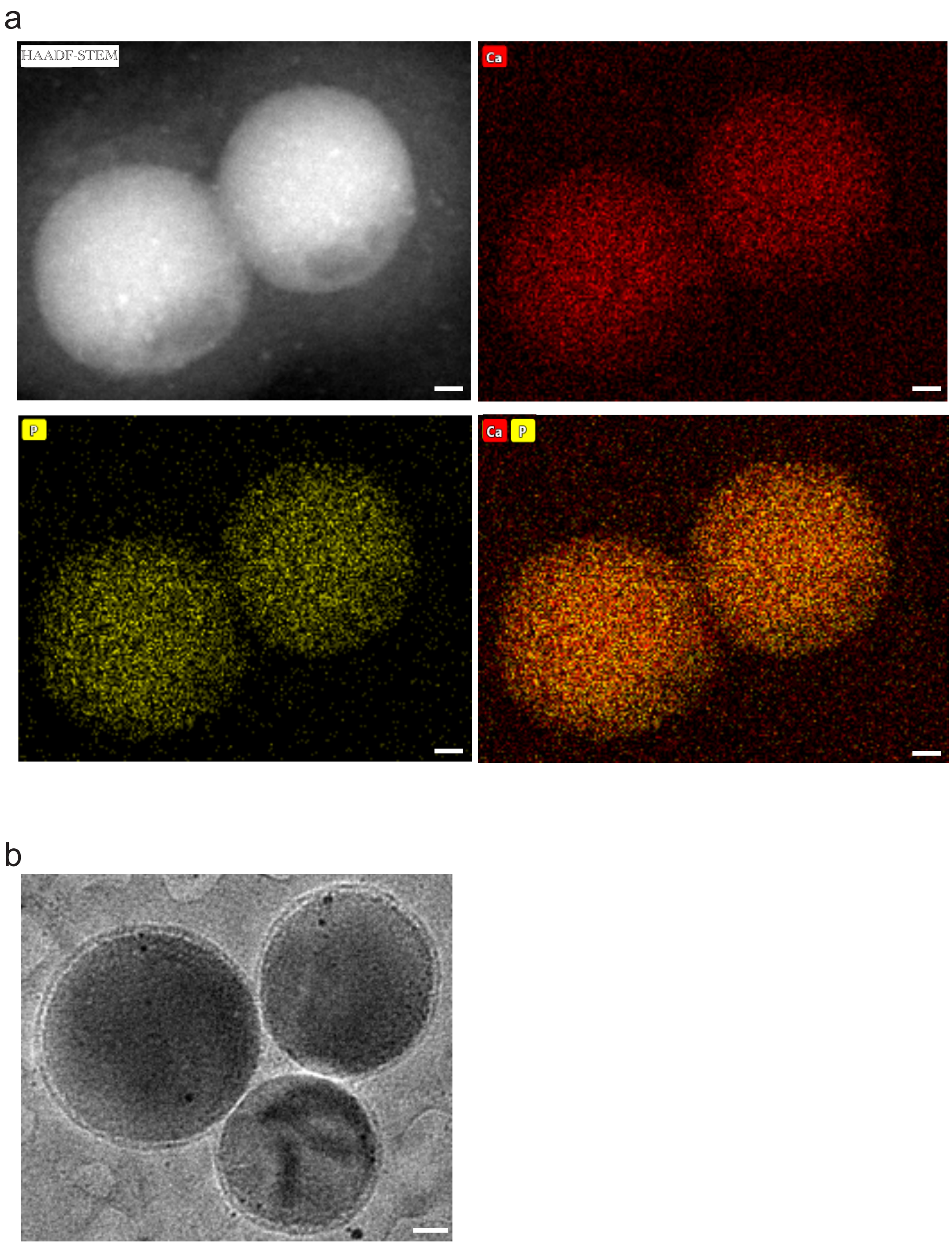


**Supporting Figure 2. Electron microscopy imaging of HAPNs and lipid/PEG-coated HAPNs (LHAPNs).** (**a**) High-angle annular dark-field scanning transmission microscopy (HAADF-STEM) imaging and STEM-energy dispersive X-ray spectroscopy (STEM-EDS) mapping of the composition of the HAPNs. (**b**) Transmission electron microscopy images of the lipid/PEG-coated HAPNs (LHAPNs) in cryo-mode. Scale bar = 10 nm.

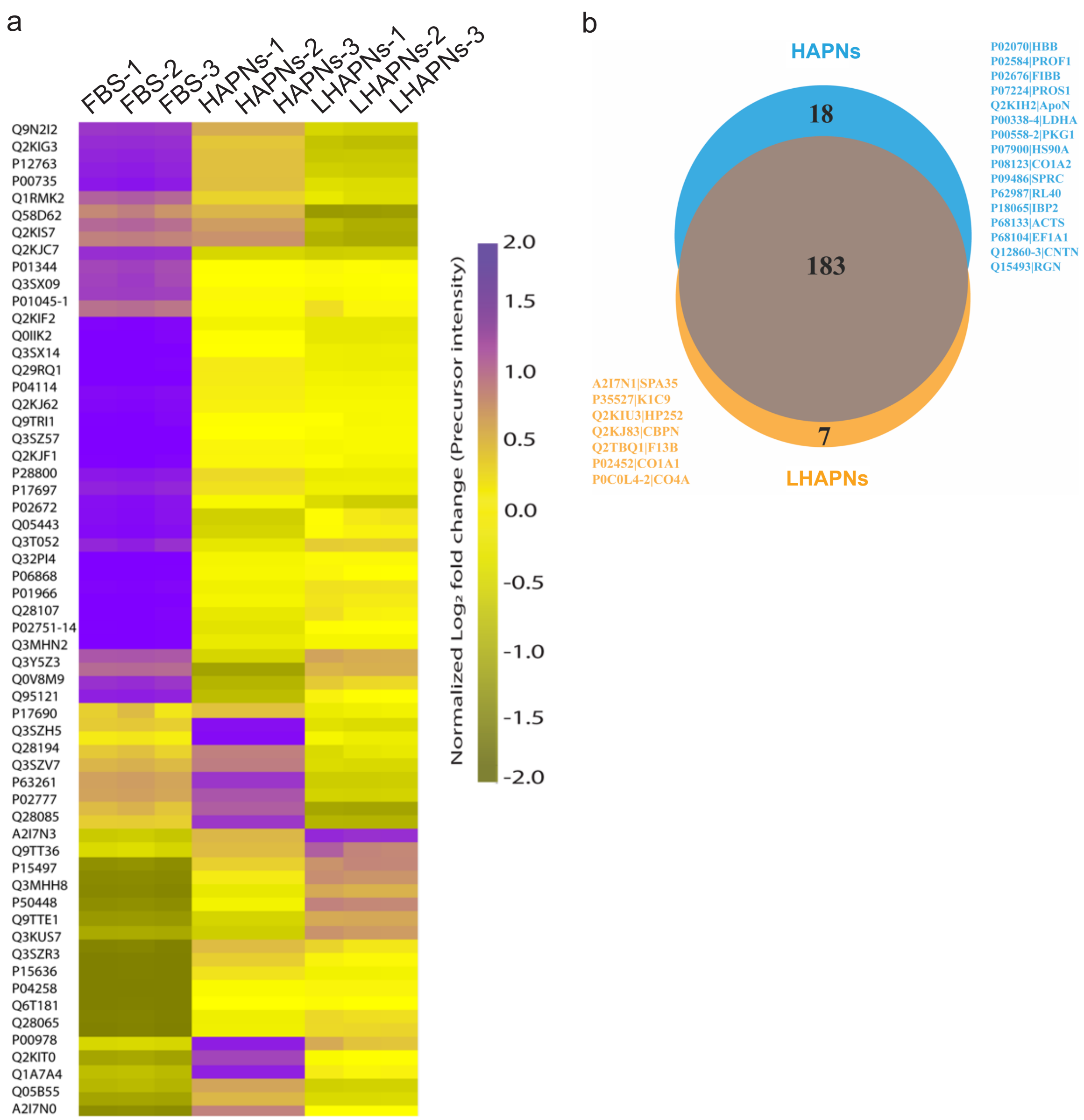


**Supporting Figure 3. Quantitative proteomic analysis of serum protein adsorption to the surface of HAPNs and LHAPNs.** (**a**) A heat map illustrating the serum proteins detected in the control fetal bovine serum (FBS) sample, as well as those adsorbed onto the surfaces of HAPNs and LHAPNs after incubation in cell culture medium (RPMI) containing 50% FBS for 72 h. The protein digests from both the control and NP samples were examined using liquid chromatography tandem mass spectrometry (LC-MS/MS), and the relative abundance of proteins was quantified through label-free quantification (LFQ).[1] As depicted in the color scale bar, the heat map uses purple to represent high LFQ intensities (log2 (LFQ)), while yellow/gold indicates lower intensities. The proteins associated with the UniProt Knowledgebase (UniProtKB)[2] accession numbers displayed in the figure are detailed in Supporting Table 2. (**b**) Venn diagram delineating adsorption of the 183 most abundant serum proteins to the surface of LHAPNs compared to HAPNs.

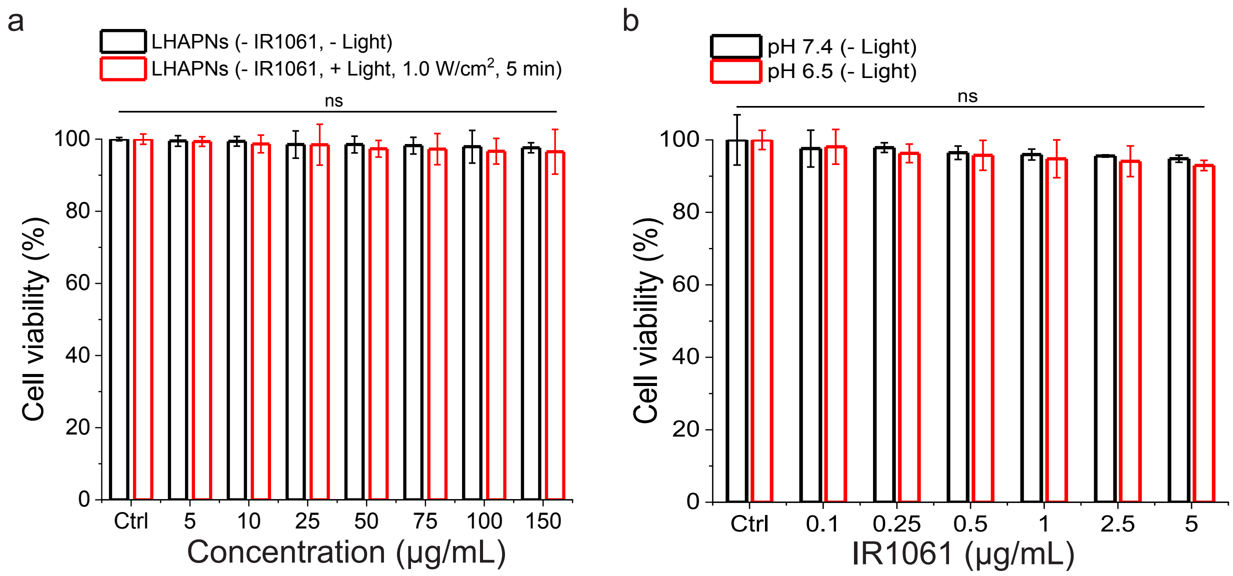


**Supporting Figure 4. Cell viability control experiments.** (**a**) Viability of MIA PaCa-2 cells treated with increasing concentrations of LHAPNs at physiological pH in the absence or presence of NIR light irradiation. (**b**) Viability of MIA PaCa-2 cells treated with increasing concentrations of LHAPNs that encapsulate the near-infrared (NIR) dye IR1061 within their pores (LHAPNIRs) at pH 7.4 or 6.5 for 48 h in the absence of NIR light irradiation. Cell viability was quantified using the MTS assay, with the % viability determined from the ratio of the absorbance of the treated cells to the control cells ($n = 3$). ns, non-significant ($P > 0.05$) for comparisons with vehicle-treated controls.

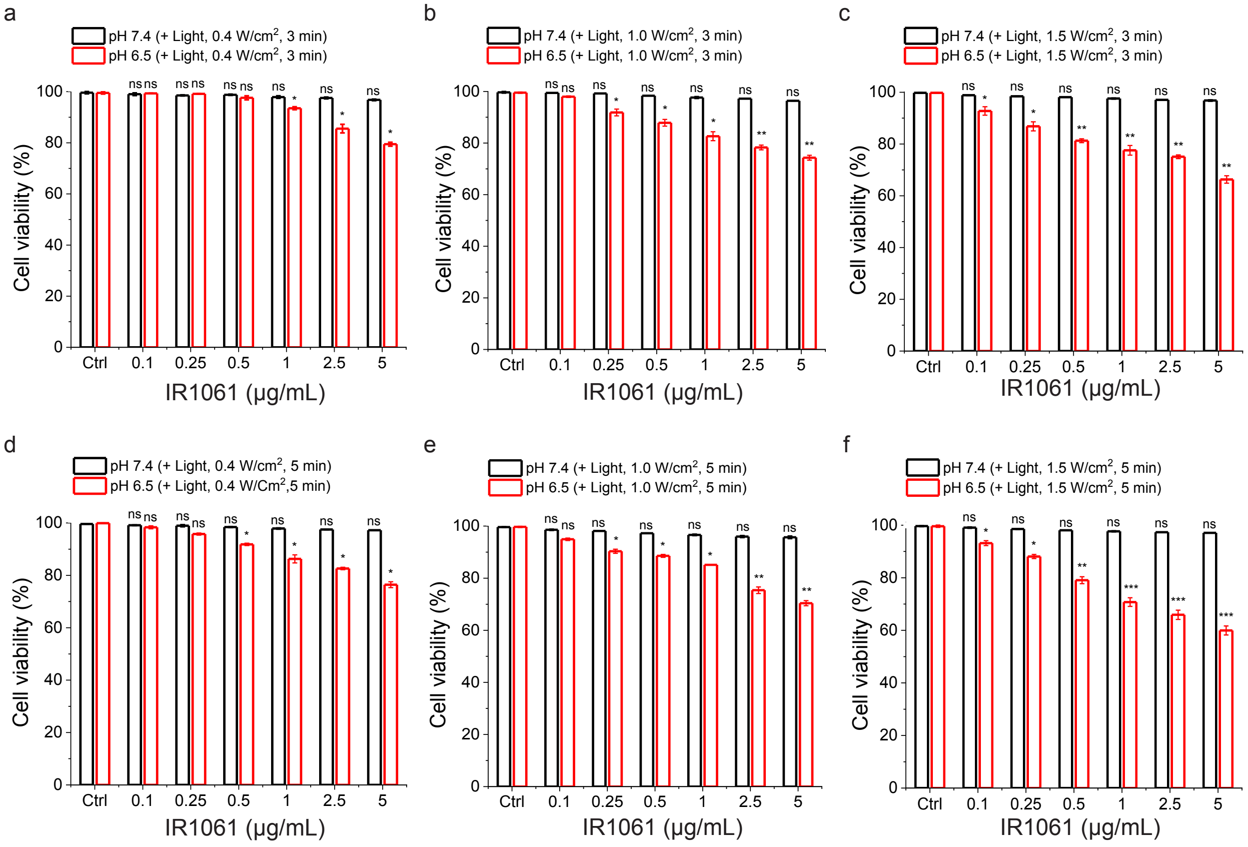


**Supporting Figure 5. NIR-light triggered cytotoxicity of the ATRAM-functionalized LHAPNIRs (ALHAPNIRs) at 24 h incubation.** Viability of MIA PaCa-2 cells treated with ALHAPNIRs (0.05–5 µg/mL IR1061) for 24 h at pHs 7.4 or 6.5 and exposed to NIR light with varying irradiation power densities (0.4–1.5 W/cm²) for 3 (**a**–**c**) or 5 (**d**–**e**) min. Cell viability was measured using the MTS assay, with the % viability determined from the ratio of the absorbance of the treated cells to the control cells ($n$ = 3). *$P$ < 0.05, **$P$ < 0.01, ***$P$ < 0.001, or non-significant (ns, $P$ > 0.05) compared with controls.

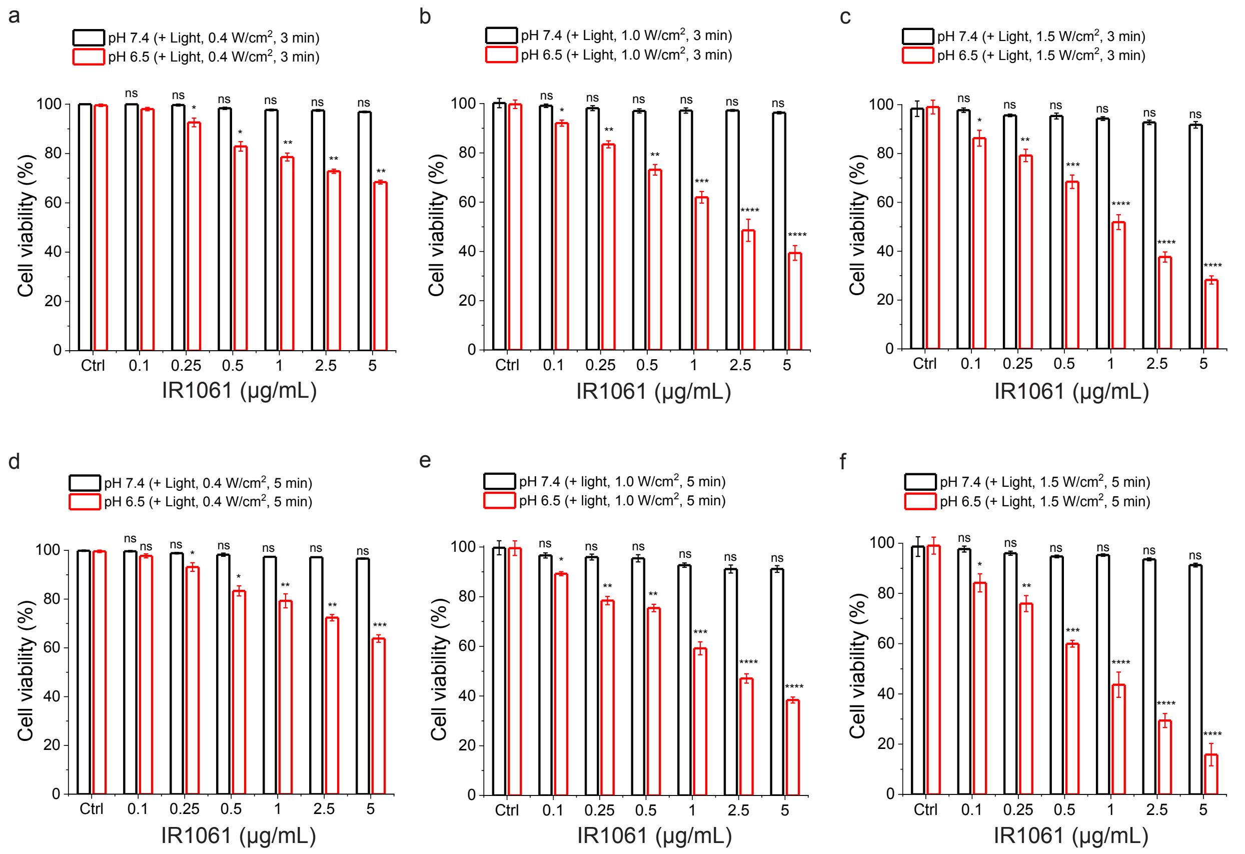


**Supporting Figure 6. NIR-light triggered cytotoxicity of ALHAPNIRs at 72 h incubation.** Viability of MIA PaCa-2 cells treated with ALHAPNIRs (0.05–5 µg/mL IR1061) for 72 h at pHs 7.4 or 6.5 and exposed to NIR light with varying irradiation power densities (0.4–1.5 W/cm²) for 3 (**a**–**c**) or 5 (**d**–**e**) min. Cell viability was measured using the MTS assay, with the % viability determined from the ratio of the absorbance of the treated cells to the control cells ($n$ = 3). *$P < 0.05$, **$P < 0.01$, ***$P < 0.001$, ****$P < 0.0001$ or non-significant (ns, $P > 0.05$) compared with controls.

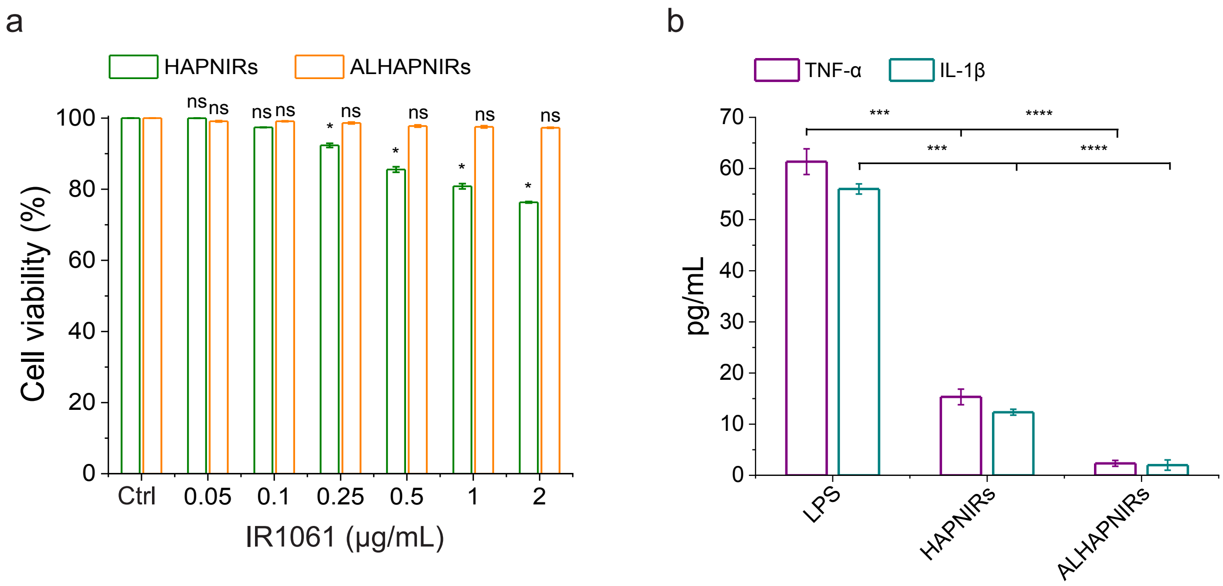


**Supporting Figure 7. Macrophage recognition and immunogenicity of the NPs.** (**a**) Cell viability of differentiated THP-1 cells treated with HAPNIRs or ALHAPNIRs for 48 h at pH 7.4. Cell viability was assessed using the MTS assay ($n = 3$). (b) Release of inflammatory cytokines, tumor necrosis factor-alpha (TNF-α) and interleukin-1 beta (IL-1β), by differentiated THP-1 cells exposed to HAPNIRs or ALHAPNIRs (0.5 μg/mL IR1061) for 24 h at pH 7.4. Cells treated with the macrophage activator lipopolysaccharide (LPS) were used as a positive control for inflammation.[3] TNF-α and IL-1β levels in the culture medium were assayed using a commercial ELISA kit ($n = 3$). *$P < 0.05$, ***$P < 0.001$, ****$P < 0.0001$ or non-significant (ns, $P > 0.05$) compared with controls.

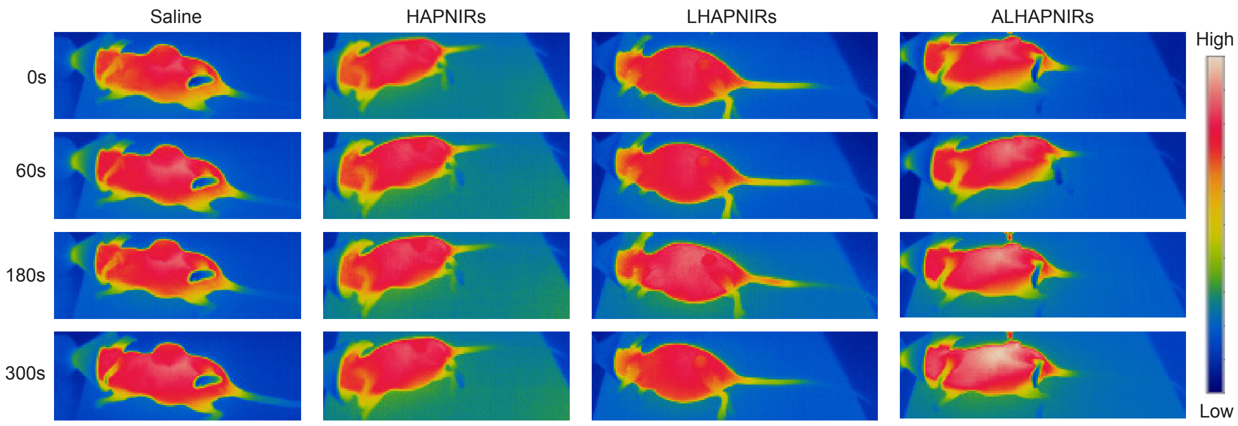


**Supporting Figure 8. ALHAPNIR-facilitated photothermal tumor imaging.** Thermal imaging of MIA PaCa-2 tumor-bearing mice upon NIR laser irradiation (1.0 W/cm$^2$, 5 min) at 8 h post *i.v.* injection with saline or HAPNIRs, LHAPNIRs or ALHAPNIRs (22.5 mg/kg nanospheres, 3.3 mg/kg IR1061) ($n$ = 4 per group).

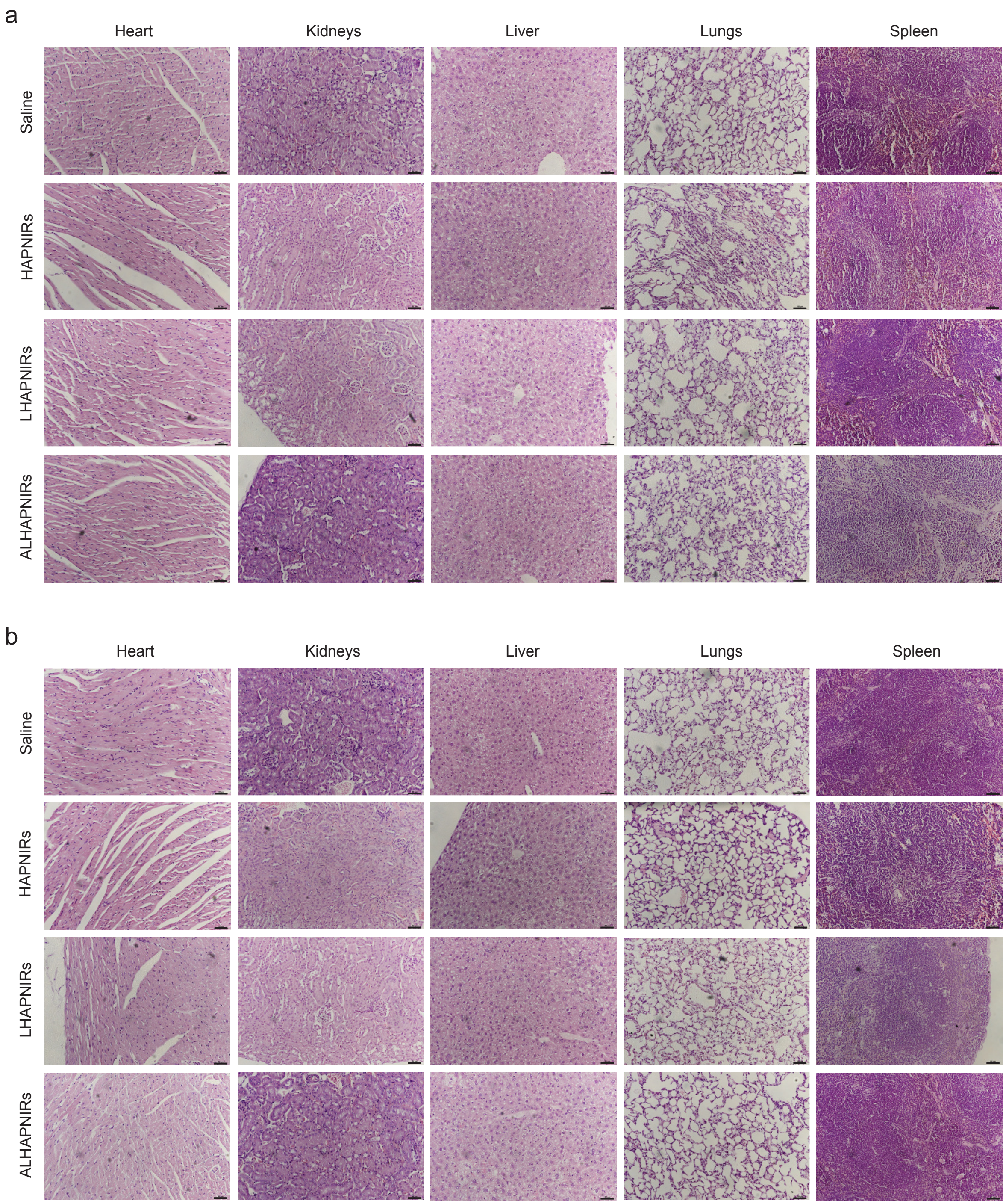


**Supporting Figure 9. Histological analysis of vital organs following treatment with the NPs.** Hematoxylin and eosin (H&E) staining of heart, kidney, liver, lung, and spleen sections from MIA PaCa-2 tumor-bearing mice after 30 days of treatment, during which mice received *i.v.* injections every two days for a total of 15 doses of saline, HAPNIRs, LHAPNIRs, or ALHAPNIRs (22.5 mg/kg NPs, 3.3 mg/kg IR1061) in the absence (**a**) or presence (**b**) of NIR laser irradiation (980 nm, 1.5 W/cm$^2$, 5 min) at 8 h post-injection. The images shown are representative of tissue sections from 4 mice per treatment group. Scale bar = 50 μm.

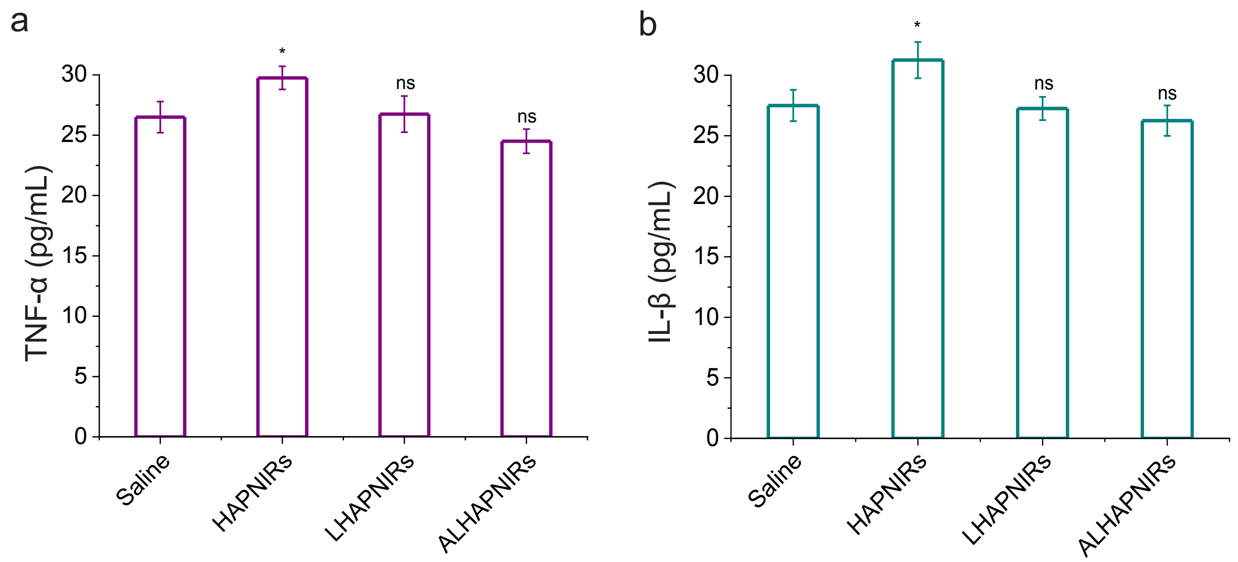


**Supporting Figure 10. Quantification of inflammatory cytokines in circulation following treatment with the NPs.** Measurement of TNF-α (**a**) and IL-1β (**b**) concentrations in serum from test mice following 30 days of treatment (*i.v.* injections administered every 2 days for a total of 15 doses) with saline and HAPNIRs, LHAPNIRs and ALHAPNIRs (22.5 mg/kg NPs, 3.3 mg/kg IR1061). TNF-α and IL-1β levels were assayed using commercial ELISA kits ($n = 4$ per group). *$P < 0.05$ or non-significant (ns, $P > 0.05$) compared with controls.

**Supporting Table 1. Summary of hydrodynamic diameters and zeta potentials of the hydroxyapatite (HAP) NPs.**

| Nanoparticles | Diameter (nm) | Zeta potential |
|---|---|---|
| HAPNs | 78 ± 10 | -6.5 |
| LHAPNs | 89 ± 8 | -13.2 |
| ALHAPNs | 90 ± 9 | -12.9 (pH 7.4)<br>+11.1 (pH 6.5) |

**Supporting Table 2. Proteins corresponding to the UniProt Knowledgebase (UniProtKB) accession numbers shown in Supporting Figure 3.**

| UniProtKB accession number | Protein |
|---|---|
| P02070 | Hemoglobin subunit beta |
| P02584 | Profilin-1 |
| P02676 | Fibrinogen beta chain |
| P07224 | Vitamin K-dependent protein S |
| Q2KIH2 | ApoN protein |
| P00338 | L-lactate dehydrogenase A chain |
| P00558 | Phosphoglycerate kinase 1 |
| P07900 | Heat shock protein HSP 90-alpha |
| P08123 | Collagen alpha-2(I) chain |
| P09486 | SPARC |
| P62987 | Ubiquitin-ribosomal protein eL40 fusion protein |
| P18065 | Insulin-like growth factor-binding protein 2 |
| P68133 | Actin, alpha skeletal muscle |
| P68104 | Elongation factor 1-alpha 1 |
| Q12860-3 | Contactin-1 |
| Q15493 | Regucalcin |

**Supporting Table 3. IR1061 dye loading capacity of the hydroxyapatite nanoparticles (HAPNs).**

| IR1061 feed ratio (to 5 mg HAPNs) | Loading capacity (wt%) |
| --- | --- |
| 0.5 | 1.0 |
| 1.5 | 6.0 |
| 2.5 | 15.0 |

The IR1061 loading capacity of HAPNs was determined as described in the Supporting Experimental Section.

### SUPPORTING EXPERIMENTAL SECTION

#### Materials

Acetone, acetonitrile, ammonium bicarbonate buffer, ammonium hydrogen phosphate, calcium nitrate tetrahydrate, chloroform, dimethylsulfoxide (DMSO), dithiothreitol (DTT), endocytosis inhibitors (amiloride, chlorpromazine, cytochalasin D and filipin), ethanol, formic acid, IR1061, isopropanol, nitric acid, phorbol 12-myristate 13-acetate (PMA), sodium acetate, sodium chloride (NaCl), sodium hydroxide (NaOH), sodium pyruvate, and trichloroacetic acid (TCA) were all purchased from Sigma Aldrich (St. Louis, MO, USA). Dead Cell Apoptosis Kit, Dulbecco's modified eagle medium (DMEM), fetal bovine serum (FBS), L-glutamine, N-2-hydroxyethylpiperazine-N-2-ethane sulfonic acid (HEPES), pencillin/streptomycin, phosphate-buffered saline (PBS), RPMI 1640, sodium pyruvate, Trypan Blue, and trypsin-EDTA were all obtained from Thermo Fisher (Waltham, Massachusetts, USA). 1,2-dipalmitoyl-sn-glycero-3-phosphocholine (DPPC), cholesterol, and the PEGylated derivative of 1,2-distearoyl-sn-glycero-PE (DSPE-$PEG_{2000}$)-maleimide were obtained from Avanti Polar Lipids Inc (Alabaster, AL, USA). Calcein AM/PI Double Staining Kit, and Human and Mouse Interleukin-1 Beta (Il-1β) and Tumor Necrosis Factor-Alpha (TNF-α) ELISA Kits were all from Elabscience (Houston, TX, USA). The CellTiter 96 Aqueous One Solution (MTS) Cell Proliferation Assay Kit was from Promega (Madison, WI, USA).

#### Synthesis of the hydroxyapatite nanoparticles

Spherical hydroxyapatite nanoparticles (HAPNs) were synthesized using a previously published method with a few modifications.[4] First, a 0.15 mol solution of calcium nitrate tetrahydrate ($Ca(NO_3)_2.4H_2O$) was prepared by dissolving 70.846 g of the compound in 400 mL Milli-Q water in a 500 mL Teflon bottle. The pH of the solution was adjusted to 7.4 using 5.0 M NaOH and this mixture was designated as solution A. In a separate 500 mL Teflon bottle, a 0.1 mol solution of ammonium dihydrogen phosphate ($NH_4.H_2PO_4$) was prepared by dissolving 26.412 g of the compound in 400 mL of Milli-Q water, and the pH was adjusted to 4.0 with 5.0 M $HNO_3$ (mixture was designated as solution B). Solution A was added to solution B at a controlled rate of 13 mL/min in a 1 L Teflon container. The pH of the resulting mixture was adjusted to pH 10.8 and the solution was stirred at room temperature for 4 h. Thereafter, the mixture was subjected to centrifugation at 10,000×g for 10 min, and the resulting precipitate was washed three times with Milli-Q water by centrifugation (10,000×g, 10 min). The obtained HAPNs were then over-dried at 100 °C for 4 h. For long-term storage, the dried HAPNs were re-dispersed in Milli-Q water and lyophilized for 48−72 h.

To load the porous HAPNs with the IR1061 fluorophore we used a slightly modified version of an established adsorption method.[5] First, 2.5–12.5 mg of IR1061 dye was dissolved in 1 mL of methanol. The resulting IR1061 solutions were then mixed with 5 mg of HAPNs at NP to dye ratios of 1:0.5, 1:1.5 and 1:2.5. Next, the resulting mixtures were sonicated for 5 min until the HAPNs were uniformly dispersed in the solution. For effective dye loading, the samples were kept on a magnetic stirrer (set to 1200 rpm) for up to 48 h at ambient temperature. Thereafter, the mixture was centrifuged at 10,000×g for 10 min to separate the IR1061-loaded HAPNs (HAPNIRs) pellet from free IR1061 in the supernatant. The HAPNIRs were then vacuum dried, washed three times with Milli-Q water to remove any residual methanol, and lyophilized for 48−72 h for further use. The mass of residual IR1061 in the isolated supernatant was determined by UV-Vis spectroscopy (Shimadzu UV-2600i Spectrophotometer) using a standard IR1061 concentration calibration curve. The IR1061 loading capacity of the HAPNs was then determined using the following formula:[6,7] loading capacity (%) = $(M_0 - M_s)/W_0) \times 100$, where $M_0$ represents the initial mass of IR1061 used in the NP loading step, $M_s$ the measured mass of IR1061 in the supernatant, and $W_0$ denotes the mass of the HAPNIRs.

Coating of the surface of the NPs with a lipid/PEG bilayer was done as previously reported.[7,8] DPPC, cholesterol, and DSPE-PEG$_{2000}$ were dissolved, at a molar ratio of 77.5:20:2.5, in a solution of chloroform-ethanol to ensure thorough mixing. After dissolution, the solvent was evaporated using an $N_2$ stream, and any remaining chloroform/ethanol traces were removed by subjecting the lipid/PEG film to a vacuum for 3 h. Following this, 5 mg of dried NPs, either without or with IR1061 (HAPNs or HAPNIRs, respectively) were dispersed in 3 mL saline solution (0.9% NaCl) and sonicated at 40 kHz for 30 s. This suspension was then deposited onto the lipid/PEG film at a NP-to-lipid ratio of 1:1.2 (*w*/*w*), and the mixture was probe sonicated for 20 min (15/15 s on/off cycle), at an output power of 30 W. Finally, the lipid/PEG-coated NPs (designated as LHAPNs or LHAPNIRs) were separated from excess lipids by centrifugation at 10,000×g for 10 min, followed by washing twice with saline and Milli-Q water.

In order to enhance tumor targeting, we modified the NPs by attaching the acidity-triggered rational membrane (ATRAM) peptide to the lipid/PEG coat. The ATRAM peptide was synthesized by Selleck Chemicals (Houston, TX, USA) using standard Fmoc solid-phase protocols, and subsequently purified in-house by reversed-phase high-performance liquid chromatography (Waters 2535 QGM HPLC), with confirmation of the peptide's purity done by mass spectrometry (Agilent 6538 QToF LC/MS). To synthesize the ATRAM-functionalized NPs, DSPE-PEG$_{2000}$-maleimide was covalently coupled to the N-terminal cysteine residue of the ATRAM peptide as previously described.[7,9] Briefly, 600 nmol ATRAM was mixed with 500 nmol DSPE-PEG$_{2000}$-maleimide in 200 μL methanol and stirred overnight under an $N_2$ atmosphere at room temperature.

Once conjugation was confirmed by mass spectrometry, the NPs – without or with the fluorophore cargo–were coated with DPPC/cholesterol/DSPE-PEG$_{2000}$-ATRAM (at a molar ratio of 77.5:20:2.5) following the aforementioned procedure. Finally, the ATRAM-functionalized NPs (ALHAPNs, ALHAPNRhBs, or ALHAPNIRs) were dialyzed against phosphate buffer and stored at 4 °C for further use.

#### Characterization of the hydroxyapatite nanoparticles

The NPs were visualized using transmission electron microscopy (TEM) on a Talos F200X (Thermo Fisher Scientific; Waltham, Massachusetts, USA) operating at 200 kV accelerating voltage. Images were acquired in brightfield TEM and high-angle annular dark-field scanning transmission microscopy (HAADF-STEM) modes. To ensure no sample damage, the maximum tolerated electron dose rate was investigated prior to imaging. Brightfield TEM imaging was performed using a spot size of 5, a gun lens of 6, and an electron dose rate of 22.0 e/Å$^2$/s, while HAADF-STEM imaging was done with a spot size of 9, gun lens 4, and a beam current < 0.2 nA. The composition of the NPs was confirmed using STEM-energy dispersive X-ray spectroscopy (STEM-EDS) mapping with a spot size of 6, gun lens 4, and an exposure time of 12.5 s/frame. To prepare the NPs for room temperature imaging, 3 μL drops of NP suspension were deposited onto freshly cleaned 300 mesh copper grids topped with an ultra-thin carbon layer (Ted Pella; Redding, CA, USA) and allowed to dry at room temperature for 2 h. For low temperature (cryogenic) imaging, the Vitrobot Mark IV System (Thermo) was used to prepare the samples. Here, a very thin layer of NP suspension is rapidly plunged into liquid ethane to embed the sample in an amorphous ice layer. Subsequently, the sample is cooled down to liquid $N_2$ temperature. Quantifoil R2/2 copper TEM grids were used as sample carrier in tandem with a Gatan cryo-transfer holder.

Specific surface area and average pore size of the NPs were determined using a 3Flux Adsorption Analyzer (Micrometrics Instruments) operated at -196 °C. Size and zeta potential of the NPs in different solutions (water; 10 mM phosphate buffer, pH 7.4; 50 mM sodium acetate buffer, pH 5.5; and RPMI 1640 cell culture medium containing 10–50% FBS, pH 7.4) were measured using a Zetasizer Nano ZS (Malvern Panalytical; Malvern, UK).

#### Quantitative proteomic analysis

To assess serum protein adsorption onto the surface of HAPNs and LHAPNs, 1 mg/mL of the NPs was incubated with cell culture medium containing 50% FBS (pH 7.4) for 72 h. Subsequently, the adsorbed serum proteins were isolated by centrifugation as previously described.[10,11] The isolated proteins were then reduced with 10 mM dithiothreitol (DTT) for 30 min at 85 °C, followed by alkylation with 25 mM iodoacetamide (IAA) for 1 h in the dark at room temperature. The proteins were precipitated using 10% trichloroacetic acid (TCA) overnight at 4 °C, after which samples were centrifuged at 16,000×g for 15 min at 4 °C. The resulting protein pellet was washed with ice-cold acetone, air-dried, and resuspended in 50 mM ammonium bicarbonate buffer to achieve the desired pH for protease activity. Digestion was performed using a trypsin/Lys-C protease mix (1:50, *w/w*) for 24 h at 37 °C. The reaction was quenched with 1 µL of formic acid, and the peptide digests were enriched through offline reversed-phase liquid chromatography (RPLC). Finally, the samples were dried and reconstituted in a 20 µL solution of 0.1% formic acid before undergoing online RPLC-tandem mass spectrometry (RPLC-MS/MS).

The RPLC-MS/MS analysis was done as previously detailed.[7] RPLC was performed on an UltiMate 3000 RSLCnano System (Dionex) equipped with a C18 column (75 µm inner diameter, 50 cm length; PepMap RSLC) maintained at 40 °C. The mobile phases used were 0.1% formic acid (solvent A) and 95% acetonitrile/0.1% formic acid (solvent B). Samples were loaded in solvent A and eluted using a gradient: starting at 0% B for 3 min, increasing to 35% B over 75 min, and then to 60% B over 10 min. A 15-min wash at 90% B prevented carryover, followed by a 20-min equilibration at 0% B. The LC system was coupled to a Thermo Q-Exactive high-field orbitrap mass spectrometer equipped with an Easy Spray ion source operating in positive ion mode. The spray voltage was set at 1.8 kV, the S-lens RF level at 35, and the ion transfer tube at 275 °C. Full scans were acquired over an m/z range of 350–1800 at a resolution of 120,000, with an AGC target set to 3E6 and maximum ion time to 50 ms. MS2 analysis was conducted in data-dependent mode to fragment the 20 most intense precursors using HCD fragmentation, with parameters set for resolution at 15,000, AGC target at 1E5, minimum AGC target at 8.0E3, intensity threshold at 4.0E5, maximum ion time at 20 ms, isolation width at 1.4 m/z, precursor charge states from 2 to 6, preferred peptide matching, dynamic exclusion for 30 seconds, and a fixed first mass of 80 m/z. Normalized collision energy for HCD was set at 32%.

For data processing, MaxQuant software (version 2.5.2.0) was used for label-free protein quantification (LFQ),[1] employing the built-in Andromeda[12] search engine with match-between-runs and iBAQ parameters. Peptide and protein false discovery rates (FDRs) were both established at 1%, while all other parameters remained at their default settings. Peptide identification was acuired using the UniProtKB_SwissProt database (https://www.uniprot.org/uniprot) with a focus on human taxonomy. Annotation and statistical analysis of the ProteinGroups files generated by MaxQuant[1] were done using Perseus (version 2.1.2.0). The data underwent filtering (removing reverse sequences and those only identified by site), with expression values log-transformed (Log2), and variations in the proteomes from the different conditions analyzed using a one-sample t-test.

#### Photothermal response

The photothermal response of LHAPNIRs was investigated by monitoring temperature increases following NIR-light irradiation of the nanoparticle samples. The samples, comprising LHAPNIRs (10–150 μg/mL) in 10 mM phosphate buffer (pH 7.4), were illuminated with an NIR laser (980 nm) at varying irradiation power densities (0–1.5 W/cm$^2$) for 5 min. For comparative purposes, samples of the LHAPNIRs (saline) and IR1061 dye alone (dissolved in saline or methanol), at the same IR1061 concentration (0.5 μg/mL), were subjected to NIR laser irradiation (980 nm, 1.5 W/cm$^2$, 0–5 min) and the temperature increases were measured. for a duration of 5–10 min, at the same concentration of IR1061 (0.5 μg/mL). To assess the photostability of the NPs, the temperature changes were monitored for 150 μg/mL LHAPNIRs over 5 consecutive NIR laser irradiation (980 nm, 1.5 W/cm$^2$, 5 min) on/off cycles. The temperatures and thermal images of the samples were recorded using a PI-640i infrared camera (Optris GmBH; Berlin, Germany).

#### Cell culture

The cell lines used in this study were acquired from the American Type Culture Collection (ATCC) and underwent authentication and testing for mycoplasma contamination (Charles River Laboratories; Margate, UK). Human Pancreatic MIA PaCa-2 cells (ATCC no. CRL-1420) were cultured in DMEM supplemented with 10% FBS, 4 mM L-glutamine, 1 mM sodium pyruvate and 1% penicillin/streptomycin at 37 °C in 5% $CO_2$. Human monocytic leukemia THP-1 cells (ATCC no. TIB-202) were cultured in RPMI 1640 medium supplemented with 10% FBS, 1 mM sodium pyruvate and 1% penicillin/streptomycin at 37 °C in 5% $CO_2$. Viability of the cells was monitored regularly during culturing using the Trypan Blue exclusion test on a Bio-Rad TC20 automated cell counter. Upon reaching ~95% confluence, the cells were harvested using 0.25% trypsin-EDTA or cell scrapers for propagation or use in the following experiments.

#### Cancer cell uptake

To assess the cellular uptake of the NPs, we utilized both confocal fluorescence microscopy and flow cytometry. For intracellular imaging, MIA PaCa-2 cells were plated at a density of $2\times10^5$ cells/well in 500 μL complete medium in 4-chambered 35 mm glass bottom Cellview cell culture dishes (Greiner Bio-One; Monroe, NC, USA) and cultured for 24 h at 37 °C in 5% $CO_2$. Thereafter, the medium was replaced with fresh medium (pH 6.5 or 7.4) containing Rhodamine B (RhB)-loaded LHAPNs or ALHAPNs (LHAPNRhBs or ALHAPNRhBs; 0.5 μg/mL RhB) and incubated for a further 4 h. Finally, the media in the chambers was replaced once again with fresh media and the cells were images on a Leica Stellaris 8 confocal microscope equipped with a 63×Plan-Apo/1.3

NA oil immersion objective with DIC capability. Images were acquired using the LASX software and analysis was performed with the Fiji image processing software e.[13]

To quantify the cellular uptake of the NPs, MIA PaCa-2 cells were cultured in 6-well plates ($2\times10^5$ cells/well) for 24 h at 37 °C in 5% $CO_2$. Subsequently, the cells were treated with ALHAPNRhBs (0.5 µg/mL RhB), at either pH 7.4 or 6.5, for 4 h. For the analysis of uptake pathways, before addition of the NPs, the cells were pre-treated for 1 h at 4 °C with a solution containing a mixture of 10 mM sodium azide and 6 mM 2-deoxy-D-glucose in serum- and glucose-free medium.[14] Alternatively, the cells were pre-treated for 30 min at 37 °C with endocytosis inhibitors (5 µM amiloride, 10 µM chlorpromazine, 5 µM filipin or 5 mM methyl-β-cyclodextrin) in serum- and glucose-free medium (pH 6.5).[14] The cells underwent a series of steps to prepare them for analysis. The cells were rinsed thrice with ice-cold PBS to remove extracellular NPs, harvested using trypsin-EDTA, followed by centrifugation (1,000×g, 5 min), and re-suspended (500 µL ice-cold PBS supplemented with 10% FBS). Data was acquired by flow cytometry (10,000 cells/sample, gated on live cells by forward/side scatter and propidium iodide (PI) exclusion) on a FACSAria III cell sorter (BD Biosciences, San Jose, CA), and analysis was performed using the Cytobank Premium software, © 2000-2025 Beckman Coulter, Inc.[15]

#### Cell viability/toxicity assays

The *in vitro* cytotoxicity of the NPs was assessed using three distinct approaches: i) the CellTiter 96 Aqueous One Solution (MTS) assay, which measures reduction of the MTS compound (3-(4,5-dimethylthiazol-2-yl)-5-(3-carboxymethoxyphenyl)-2-(4-sulfophenyl)-2H-tetrazolium, inner salt) into a soluble formazan product by intracellular dehydrogenases in viable cells;[16,17] ii) calcein AM/propidium iodide (PI) staining, where calcein AM, a cell-permeable non-fluorescent dye, is converted into a fluorescent calcein by esterases in living cells, while the membrane-impermeant PI, a red-fluorescent dye that binds to nucleic acids, serves as a counterstain;[18,19] and iii) the Dead Cell Apoptosis assay, which uses Alexa 488-conjugated annexin V to detect exposed phosphatidylserine in cells undergoing apoptosis, and PI to distinguish between apoptotic and necrotic cells by assessing membrane integrity.[20]

The MTS assay was done according to a published protocol.[6,7,21,22] MIA PaCa-2 cells were seeded at a density of $5\times10^3$ cells/well in 100 µL complete medium in 96-well plates and cultured (37 °C/5% $CO_2$) for 24 h. Thereafter, the medium was replaced with fresh serum-free medium (at pH 7.4 or 6.5) containing the specified concentrations of the NPs. The cells were incubated for 6 h before the cell culture medium was replaced with fresh medium to remove extracellular NPs, and the cells were exposed to NIR (980 nm) laser light at the specified irradiation power densities and durations. After culturing for an additional 18–66 h (for total NP incubation times of 24–72 h), 20 μL MTS reagent was added to each well, and the plates were incubated for 4 h. The absorbance of the formazan product (λ = 490 nm) of MTS reduction was measured using a Cytation 5 Multi-Mode Microplate Reader (BioTek; Winooski, VT, USA). Cells treated with vehicle alone served as control, while wells with medium alone were used as a blank. Cell viability was determined from the ratio of formazan absorbance of the treated cells to the controls.

Calcein AM/PI double staining was performed as previously described.[7,18] Briefly, MIA PaCa-2 cells were seeded at a density of $2\times10^5$ cells/well in 500 μL complete medium in 4-chambered 35 mm glass bottom Cellview culture dishes. After culturing (37 °C, 5% $CO_2$) for 24 h, the cells were treated with ALHAPNIRs (0.5 μg/mL IR1061 in fresh serum-free medium, pH 6.5) for 12 h. Thereafter, the cell culture medium was replaced with fresh medium to remove extracellular NPs, with or without subsequent exposure to NIR laser irradiation (980 nm, 1.0 W/cm$^2$, 5 min). Next, the cells were stained with 2 μM calcein AM and 1.5 μM PI for 30 min, and imaged on a Leica Stellaris 8 confocal microscope equipped with a 63×Plan-Apo/1.3 NA oil immersion objective with DIC capability. Images were acquired using LASX and analyzed with Fiji.

The Dead Cell Apoptosis assay carried out as previously reported.[6,7,21] Briefly, MIA PaCa-2 cells were seeded ($1\times10^6$ cells/well) in complete medium in 6-well plates and cultured (37 °C, 5% $CO_2$) for 24 h. The medium was then replaced with fresh serum-free medium containing ALHAPNIRs (0.5 μg/mL IR1061) for 48 h at pH 6.5, with or without exposure to NIR laser light (1.5 W/cm$^2$, 5 min) at the 6 h mark. Subsequently, the cells were washed twice with ice-cold PBS, harvested by trypsinization, centrifuged (1,000×*g*, 5 min) and re-suspended in 1× annexin V-binding buffer (10 mM HEPES, 140 mM NaCl, 2.5 mM $CaCl_2$, pH 7.4). Finally, the cells were stained with 5 μL FITC-conjugated annexin V and 1 μg/mL PI for 30 min in the dark at ambient temperature. Data was acquired by flow cytometry (30,000 cells/sample) and analyzed using the Cytobank Premium software, © 2000-2025 Beckman Coulter, Inc.[15]

#### Macrophage toxicity and immunogenicity

For viability analysis, differentiated THP-1 cells were cultured in standard 96-well plates ($5\times10^3$ cells/well) for 24 h at pH 7.4. The medium was then replaced with fresh medium containing LHAPNIRs or ALHAPNIRs (0.05–2 μg/mL IR1061) and the cells were incubated for a further 48 h. Thereafter, cell viability was assessed using the MTS assay, with the % viability determined from the ratio of the absorbance of the treated cells to the control cells.

To assess the release of inflammatory cytokines, tumor necrosis factor-alpha (TNF-α), and interleukin-1 beta (IL-1β), that are primarily generated by macrophages/monocytes during acute inflammation,[23] differentiated THP-1 cells were cultured in standard 96-well plates ($2\times10^4$

cells/well in 100 μL medium) for 24 h at pH 7.4. Subsequently, the medium was replaced with fresh medium containing LHAPNIRs and ALHAPNIRs (0.5 μg/mL IR1061), and the cells were incubated for a further 24 h. The culture medium was then assayed for TNF-α and IL-1β secretion using commercial enzyme-linked immunosorbent assay (ELISA) kits. Cells treated with the macrophage activator lipopolysaccharide (LPS)[3] served as positive controls, while vehicle-treated treated cells were used as negative controls. The total levels of TNF-α and IL-1β were determined by measuring absorbance (λ = 450 nm) using a Cytation 5 Multi-Mode Microplate Reader (BioTek; Winooski, VT, USA) using a standard TNF-α concentration calibration curve.

#### *In vivo* experiments

All animal experiments were done in accordance with protocols approved by the NYU Abu Dhabi Institutional Animal Care and Use Committee (NYUAD-IACUC; Protocol No. 21-0005), and adhered to established animal care and use guidelines.[24] Experiments were performed on 6–8 week-old female BalbC nude mice (The Jackson Laboratory; Bar Harbor, ME, USA) that were maintained in air-filtered cages (20 °C, 50% humidity, 12 h light/dark cycle) and fed normal chow (Research Diets, New Brunswick, NJ) in the NYU Abu Dhabi Vivarium Facility.

Tumors were induced by injecting $2\times10^5$ viable MIA PaCa-2 cells subcutaneously into the right flank of the mice. Precise measurements of tumor dimensions were taken using high-precision calipers (Thermo Fisher), and tumor volume was calculated as follows: tumor volume ($mm^3$) = $(W^2 \times L) / 2$, where W and L represent the tumor's width and length in mm, respectively. Experiments were initiated once the tumor volume reached 25 $mm^3$. Daily monitoring of the mice was conducted, and euthanasia was performed upon reaching the predefined tumor volume threshold established by NYUAD-IACUC guidelines.

#### Biodistribution and tumor accumulation

The *in vivo* biodistribution and accumulation of the NPs was assessed using thermal and fluorescence. To conduct *in vivo* thermal imaging, mice bearing MIA PaCa-2 tumors received a single intravenous injection of either saline or HAPNIRs, LHAPNIRs or ALHAPNIRs (22.5 mg/kg NPs, 3.3 mg/kg IR1061) ($n$ = 4 per group). At 8 h post-injection, the tumors were irradiated with NIR laser light (1.0 W/$cm^2$, 5 min) and thermal images were acquired using an Optris PI-640i infrared camera.

For *in vivo* fluorescence imaging, MIA PaCa-2 tumor-bearing mice received a single intravenous injection of either saline or DiD-loaded NPs (HAPNDiDs, LHAPNDiDs or ALHAPNDiDs; (5 mg/kg nanoparticles, 0.3 mg/kg DiD) ($n$ = 4 per group). The mice were then anesthetized and imaged using an *in vivo* optical imaging system (Perkin Elmer IVIS Spectrum;

Shelton, CT, USA). Imaging parameters were set as follows: $\lambda_{ex/em}$ = 605–650/650–720 nm; a binning factor of 8; an f/stop of 2; a field of view set to C; and an automatic exposure time of 0.5–1 s. Spectral unmixing was performed using the Living Image software to separate the probe signal from tissue autofluorescence, which was established using tumors of untreated control mice. Images were analyzed by drawing regions of interest (ROIs) around the tumor of each animal from the unmixed images. Average radiance efficiency (normalized data) was used for quantification. *In vivo* images were acquired at different time points (0–72 h) following *i.v.* injection. Finally, the mice were euthanized, and the tumors and vital organs were isolated for *ex vivo* fluorescence imaging to determine the biodistribution of the DiD-loaded NPs.

#### Tumor growth inhibition

MIA PaCa-2-tumor bearing mice were randomly assigned to one of four treatment groups ($n$ = 16 per treatment group), which were injected intravenously with saline, IR1061 (3.3 mg/kg), or LHAPNIRs or ALHAPNIRs (22.5 mg/kg NPs, 3.3 mg/kg IR1061). This concentration of IR1061 was chosen due to its effectiveness in studies of PTT-mediated tumor growth inhibition.[25] Injections were done every other day, for a total of 15 doses, with the first day of treatment marked as day 0. Within each treatment group, half of the mice were subjected to NIR laser irradiation (980 nm, 1.5 W/cm$^2$, 5 min) at 8 h post-injection. Tumor volume and bodyweight were recorded every other day for the duration of treatment. After the 30 days of treatment, mice ($n$ = 4 per group) were euthanized to determine the tumor mass and for histological analysis of the tumors and vital organs (heart, kidneys, liver, lungs, and spleen).

Histological analysis was done as previously described.[6,7] Briefly, the isolated tissue samples were fixed in 10% formalin, embedded in paraffin, and sectioned into 4-µm slices using a Leica RM2265 microtome. The resulting tissue sections were then dewaxed and stained with hematoxylin and eosin (H&E) using standard procedures.[26,27] Finally, the tissue sections were imaged using a NIKON LV100 upright microscope, and the acquired images were processed using the ECLIPSE LV software.

For quantification of inflammatory cytokines in serum, mice were injected intravenously (every 2 days for a total of 15 doses) with saline, HAPNIRs, LHAPNIRs or ALHAPNIRs (22.5 mg/kg NPs, 3.3 mg/kg IR1061; $n$ = 4 per group). At the end of the 30 days of treatment, the mice were sacrificed and 0.75 mL blood was drawn from the abdominal vena cava. Subsequently, the blood was centrifuged (1,000×$g$, 10 min) and the serum was carefully collected using a fine-bore pipette. Finally, concentrations of TNF-α and IL-1β in the serum samples were quantified by ELISA using commercial kits.

#### Statistical analysis

To ensure unbiased results, different investigators independently, and in a blinded manner, performed the different parts of experiments (i.e. treatment, data collection, and data analysis). Sample sizes for the *in vivo* studies were calculated using power analysis based on NYUAD-IACUC Protocol No. 21-0005. Error bars in this study represent the mean ± standard deviation from at least three independent replicates (i.e. $n \geq 3$). Statistical analysis was done using GraphPad Prism (version 10). Unless otherwise stated, an unpaired t-test was used to determine statistical significance between two groups, whereas for comparisons involving three or more groups, one-way analysis of variance (ANOVA) followed by Dunnett's or Tukey's post hoc test was applied. A $P$-value of $< 0.05$ was considered statistically significant.
